# Diversity Across the Pancreatic Ductal Adenocarcinoma Disease Spectrum Revealed by Network-Anchored Functional Genomics

**DOI:** 10.1101/2020.09.17.302034

**Authors:** Johnathon L. Rose, Sanjana Srinivasan, Wantong Yao, Sahil Seth, Michael Peoples, Annette Machado, Chieh-Yuan Li, I-Lin Ho, Jaewon J. Lee, Paola A. Guerrero, Eiru Kim, Mustafa Syed, Joseph R. Daniele, Angela Deem, Michael Kim, Christopher A. Bristow, Eugene J. Koay, Giannicola Genovese, Andrea Viale, Timothy P. Heffernan, Anirban Maitra, Traver Hart, Alessandro Carugo, Giulio F. Draetta

## Abstract

Cancers are highly complex ecosystems composed of molecularly distinct sub-populations of tumor cells, each exhibiting a unique spectrum of genetic features and phenotypes, and embedded within a complex organ context. To substantially improve clinical outcomes, there is a need to comprehensively define inter- and intra-tumor phenotypic diversity, as well as to understand the genetic dependencies that underlie discrete molecular subpopulations. To this end, we integrated CRISPR-based co-dependency annotations with a tissue-specific co-expression network developed from patient-derived models to establish CoDEX, a framework to quantitatively associate gene-cluster patterns with genetic vulnerabilities in pancreatic ductal adenocarcinoma (PDAC). Using CoDEX, we defined multiple prominent anticorrelated gene-cluster signatures and specific pathway dependencies, both across genetically distinct PDAC models and intratumorally at the single-cell level. Of these, one differential signature recapitulated the characteristics of classical and basal-like PDAC molecular subtypes on a continuous scale. Anchoring genetic dependencies identified through functional genomics within the gene-cluster signature defined fundamental vulnerabilities associated with transcriptomic signatures of PDAC subtypes. Subtype-associated dependencies were validated by feature-barcoded CRISPR knockout of prioritized basal-like-associated genetic vulnerabilities (*SMAD4*, *ILK*, and *ZEB1*) followed by scRNAseq in multiple PDAC models. Silencing of these genes resulted in a significant and directional clonal shift toward the classical-like signature of more indolent tumors. These results validate CoDEX as a novel, quantitative approach to identify specific genetic dependencies within defined molecular contexts that may guide clinical positioning of targeted therapeutics.

## INTRODUCTION

Pancreatic ductal adenocarcinoma (PDAC) is the third leading cause of cancer-related death in the United States, with a 5-year survival rate of 8% and a median survival of <11 months^1,2^. Despite continual attempts to manage this disease with targeted drugs and immunotherapy, a vast majority of PDAC tumors are recalcitrant to therapeutic interventions. This is partially due to the dominance of *KRAS* gain-of-function mutations (90% of tumors), and frequent loss-of-function genetic alterations in the well-known epithelial tumor suppressors, *TP53* (64%), *SMAD4* (23%), and *CDKN2A* (17%), which make this disease resistant to RTK pathways inhibitors, inducers of apoptosis, and other drugs. Beyond these dominant genetic alterations, mutation-based diversity in PDAC consists of other mutations present at a significantly lower frequency (<10%)^3,4^. This low mutation load is likely one factor that limits the efficacy of immunotherapy, as checkpoint inhibitor monotherapies have shown little success in unselected patients^5^.

The fundamental functional characteristics, or “hallmarks”, of tumor cells, predict that interfering with functions such as cell division, cell motility, cell energetics, etc., should profoundly inhibit tumor growth and progression. Yet, redundancy in essential pathways and adaptation often result in selective enrichment of drug-resistant tumor cells and disease progression and relapse. Recent efforts in PDAC molecular subtyping have better characterized disease heterogeneity using transcriptomic signatures associated with clinical features, which can be used to define multiple PDAC subtypes^6–9^. Of these, the most widely accepted classification distinguishes between “classical” and “basal-like” tumor subtypes^7^. While these subtypes possess prognostic relevance, with basal-like tumors exhibiting the poorer prognosis, their utility to predict patient response to specific targeted therapeutics has not yet been realized. Moreover, clonal and sub-clonal evolution as well as therapeutic intervention can result in molecular signatures that change the original subtype classification, highlighting the relevance of intratumoral molecular and functional heterogeneity that underlies high-level subtype groupings^10,11^.

We have developed a systematic approach to correlate the status of pathway activation in a tumor with its response to genetic suppression of individual gene functions. To associate genetic drivers with clinically predictive transcriptomic signatures, we developed CoDEX, a dedicated PDAC co-expression network to anchor dependencies within annotated cluster signatures. CoDEX represents the integration of CRISPR-based co-dependency annotation with a tumor-specific co-expression network that we derived from a curated cohort of patient-derived xenografts (PDXs), and it allows us to quantitatively delineate context-specific vulnerabilities within high-resolution maps of transcriptomic diversity.

To develop CoDEX, we first refined and annotated the PDX PDAC co-expression network into 31 biologically defined gene clusters. Then, using prominent anticorrelated cluster signatures derived from the co-expression network, we defined PDAC classical and basal-like subtypes on a continuous scale and quantified the transitory nature of the transcriptional signature underlying these molecular subtypes, which defined a group that exhibited a “quasi-basal” signature. We captured the subtype-associated signatures, observed in PDX models, across TCGA tumor samples, and also at the single-cell level in sequenced PDX-derived cell lines and human PDAC primary tumor samples. Finally, leveraging our previously optimized *in vivo* genetic interference screening platform^12,13^, we used a customized CRISPR library to expand on interacting nodes identified through a protein-protein interaction network, which identified a spectrum of interconnected dependencies characteristic of each PDX model tested.

CoDEX is a novel approach to establish context specificity of individual gene targets within PDAC molecular subtypes. This bears translational significance for several reasons. First, gene centrality serves as a quantitative method to prioritize cluster-representative genes, with centrality and direct co-expression jointly defining ideal biomarkers for subtype-specific dependencies. Second, by anchoring dependencies identified through CRISPR screening within the co-expression network, we are able to uncover anti-correlative cluster signatures that inform on the molecular context underlying each molecular subtype. Fundamental dependencies associated with the basal-like subtype were validated intratumorally using feature barcoding and single-cell RNA sequencing (scRNAseq), which directly supported the functional relevance of specific genetic perturbations both at the level of sub-clonal composition and network-defined transcriptomic signatures. Third, CoDEX annotation of bulk tumors uncovers an opportunity to selectively eradicate dominant sub-populations and explore transcriptomic heterogeneity as a metric to characterize tumor response to perturbation, as opposed to tumor size alone.

In sum, this work describes the application of a novel, quantitative methodology to characterize and stratify genetic targets on transcriptomic signature patterns, which represents an important advancement toward the development of subtype-specific targeted therapies in PDAC.

## RESULTS

### Defining Transcriptomic Diversity in PDAC through the Construction and Annotation of a PDX-based Co-Expression Network

To assess the diversity of transcriptomic signatures among PDAC tumors, we selected early passage patient-derived PDAC xenografts, which maintain the cellular heterogeneity of tumor lesions while reducing the contribution of the stromal components prevalent in these tumors^14^. We performed whole-transcriptome sequencing on a set of 48 PDAC PDX tumors curated at MD Anderson (Figure 1A). Within this cohort, we recognized a level of transcriptional diversity consistent with the previously defined Moffitt classification status and distribution of classical and basal-like pancreatic adenocarcinoma (Extended Data Figure 1A). In addition, we conducted whole-exome sequencing, which confirmed mutation frequencies comparable to those reported by the TCGA^4^ (Figure 1B).

**Figure 1:**
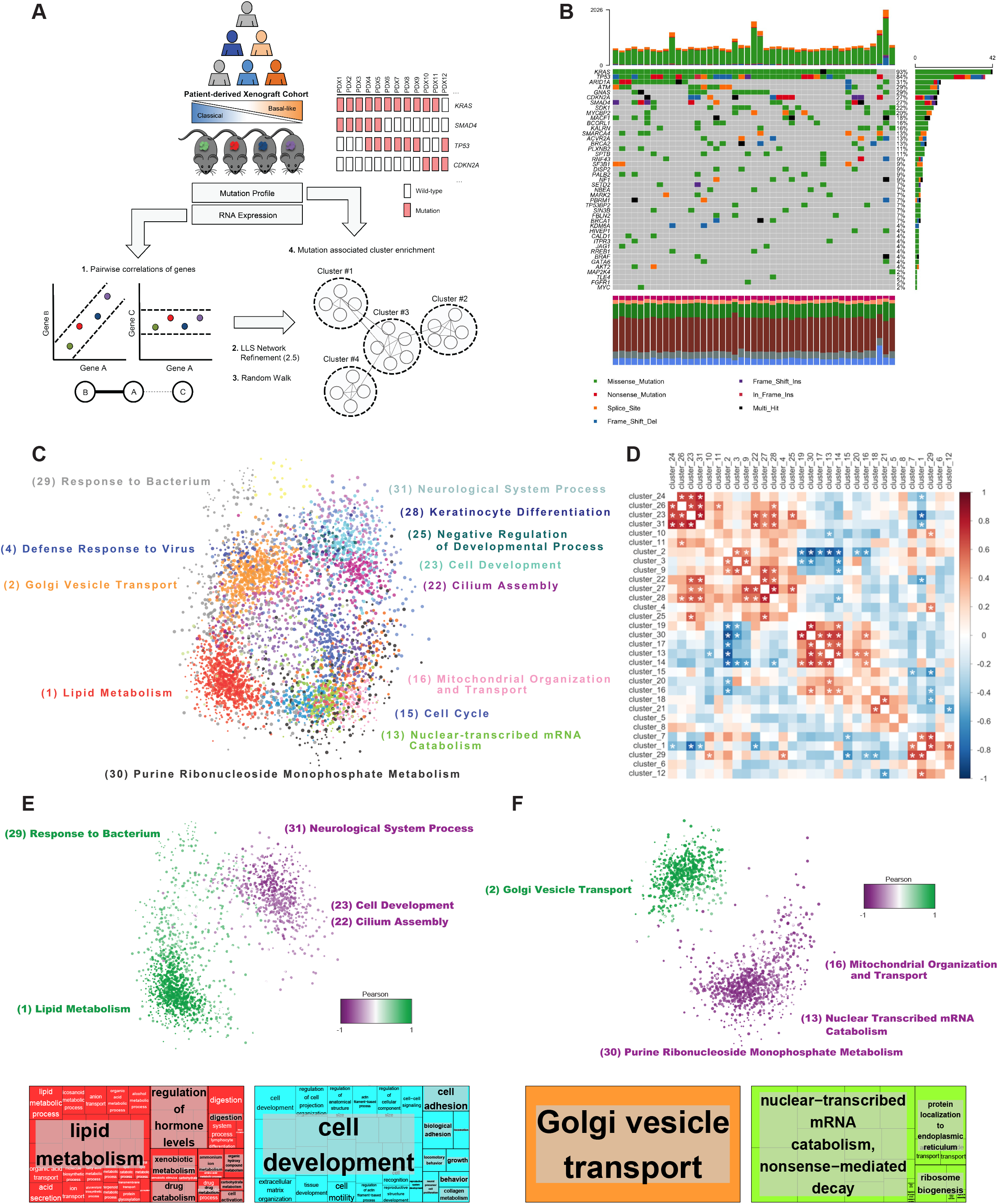
Defining signaling diversity in PDAC through the construction and annotation of a PDX-based co-expression network. (**a**) Overview outlining the development of the PDAC co-expression network from a diverse and representative PDAC patient-derived xenograft (PDX) cohort of 48 models, and subsequent integration of mutation data for cluster enrichment analysis. (**b**) Oncoplot of common PDAC-associated mutations in the 48-model PDX cohort displays mutation frequencies similar to previous publications. (**c**) Force-directed layout of the PCEN. Gene clusters, defined by Gene Ontology (GO), visually highlight nodes with a minimum of 12 edges. (**d**) Heat map depicting Pearson correlations between cluster centroid scores across 48 PDX models. Starred correlations represent an adjusted P value < 0.05. (**e**) Force-directed layout of the PCEN (top) and matching GO hierarchical treemaps (bottom) of prominent anti-correlative cluster centroid trends across the PDAC co-expression network highlighting Cluster 1 (Lipid metabolism) vs. Cluster 23 (Cell development). (**f**) Force-directed layout of the PCEN (top) and matching GO hierarchical treemaps (bottom) of prominent anti-correlative cluster centroid trends across PDAC co-expression network highlighting Cluster 2 (Golgi vesicle transport) vs. Cluster 13 (mRNA catabolism).

To characterize our cohort, we calculated global pairwise Pearson correlations of genes with variable expression to quantify concordant patterns of transcriptomic diversity across our models. These correlations were further pruned to prioritize biologically relevant gene pairs^15^, and from the resulting 103,000 correlations of 7,828 genes, we established a PDAC co-expression network (Figure 1A and Extended Data Figure 1B). Using InfoMap^16^, a community detection tool, we divided the co-expression network into 31 clusters, which were further genome ontology (GO) annotated (Figure 1C). We measured PDAC diversity by applying a dimensional reduction approach on our 31 defined clusters. Specifically, we quantified the mean expression of each cluster, or centroid score, on a tumor-by-tumor basis, for all 48 PDX models in the PDAC cohort (Figure 1D). To determine if the mutational background of a model was significantly associated with cluster enrichment across the PDX cohort, we applied UNCOVER^17^, a method to identify complementary patterns of mutation enrichment across groups.

We observed general patterns of anti-correlative clusters across the PDAC PDX cohort that were reflected in cluster positioning and subsequent cross-cluster connectivity. The most significant anticorrelated signatures were identified in clusters predominantly localized to adjacent ends of the force-directed layout (Figure 1D – F), and the non-overlap of these adjacent anti-correlated cluster trends implicates multiple distinct molecular signaling contexts that represent the immense diversity across the PDAC disease spectrum. The top anti-correlative signatures were quantified between two opposing clusters: Cluster 1 vs. Cluster 23, respectively enriched for lipid metabolism vs. cell development; and Cluster 2 vs. Cluster 13, respectively enriched for Golgi-vesicle transport vs. nuclear-transcribed mRNA catabolism (Figure 1E – F). Additionally, the PDAC co-expression network also highlighted GO annotated clusters critical for proliferating tumor cells; specifically, Cluster 15, cell cycle, and Cluster 16, mitochondrial transport and organization (Figure 1C and Extended Data Figure 1D). These findings reveal that, along with 31 unique gene clusters, two distinct anti-correlated cluster signatures contribute to PDAC tumor diversity and have the potential to provide context for tumor cell-intrinsic vulnerabilities.

Interestingly, we identified a limited cluster-mutation association between *ACVR2A*, *RREB1*, and *MARK2* mutations and tumors with significant enrichment in Cluster 21 (Extended Data Figure 1C). PDAC-associated loss-of-function mutations in *RREB1*, which encodes a zinc finger transcription factor that binds to RAS-responsive elements, have been reported in the TCGA PDAC cohort^4^. *RREB1* is a positive regulator of the zinc transporter, ZIP3, with loss-of-function playing a potential role in limiting zinc uptake and shielding developing tumors from the cytotoxic effects of high cellular zinc concentrations^18^. *RREB1* has also been described as a KRAS-regulated SMAD co-factor involved in driving the expression of epithelial-to-mesenchymal (EMT) transcription factors^19^. While the significance of mutations in the serine-threonine kinases, *MARK2* and *ACVR2A*, is relatively poorly understood in PDAC, dysregulation in these genes could implicate the regulation of epithelial polarity and downstream SMAD-associated signaling, respectively^20–22^. Associating co-expressed and annotated gene clusters with these less frequent mutations in PDAC could aid in illuminating the molecular signaling, and potential therapeutic avenues, underlying these genomic alterations.

### *In vitro* and *in vivo* screening of essential signaling pathways in PDAC

We selected four PDXs out of those used to establish the co-expression network based on the presence of mutations representative of PDAC (Extended Figure 2B and 2C), and we employed early passage cell lines (PDX lines) of these models for genetic screening. We applied a stepwise custom functional genomics platform for screening Moffitt-defined classical (PATC69) and basal-like (PATC124, PATC53, and PATC153) PDX lines in parallel *in vivo* and *in vitro* (Figure 2A and Extended Data Figure 2A). Small, customized lentiviral libraries were used to ensure maintenance of library complexity *in vivo*, based on tumor-initiating cell frequency assessment^12^.

**Figure 2.**
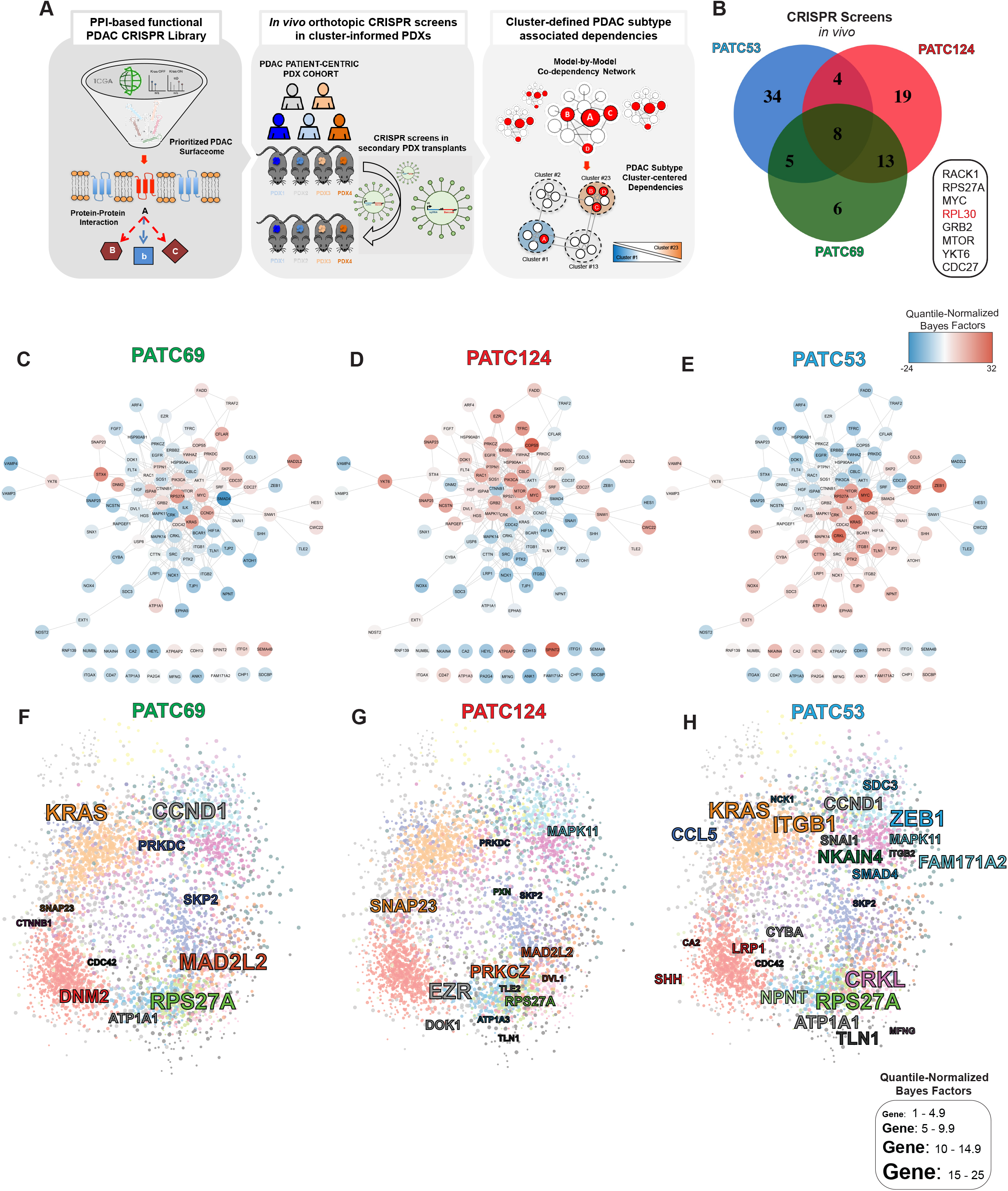
Informed CRISPR characterization of PDX models identifies co-expression network cluster-associated functional diversity within the PDAC cohort. (**a**) Overview of our prioritized *in vivo* functional genomics platform, whereby a custom PDAC-prioritized RNAi library was used to screen a functional set of proteins largely localized at the cell surface. Functionalized surface proteins were then expanded upon using a protein-protein interaction (PPI) network, which was further characterized using a custom CRISPR library and integrated into the PCEN. (**b**) Venn diagram of functional targets derived from *in vivo* orthotopic CRISPR screening of PDX lines using a quantile-normalized Bayes factor (BF) > 1 (**c - e**) CRISPR screening results from PATC69 (c), PATC124 (d), and PATC53 (e) PDX lines overlaid onto a merged PPI force-directed diagram. The BF of each gene indicates the degree of vulnerability of the gene, with BF > 1 indicating an essential gene. (**f - h**) PCEN anchoring of *in vivo* functional dependencies, represented as quantile-normalized Bayes Factors, to inform on cluster-associated vulnerability context in PATC69 (f) PATC124 (g), and PATC53 (h) PDX lines.

To perform *in vivo* and *in vitro* RNAi screens, we designed a surface proteins-targeting library (Figure 2A and Extended Table 1A) based on evidence of differential expression in pancreatic tumors compared to matched-normal tissue, copy number versus RNA expression correlation trends from the TCGA PDAC dataset, and SILAC screening for mutant *Kras* dependency (Extended Data Table 1A)^4,23–27^. We used redundant shRNA activity (RSA) to statistically score and identify candidate protein hits whose targeting shRNA were selectively depleted in the screens (Extended Data Figure 2D – G). Through these initial RNAi screens, we functionally annotated a wide range of PDAC-associated dependencies at the cell surface and uncovered oncogenic signaling diversity across the PDX lines (Extended Data Figure 2).

We subsequently built a custom sgRNA library that re-annotated and expanded upon the RNAi-identified protein surface dependencies (See Methods) across all of our PDX models (Figure 2A, Extended Data Figure 2H). Using a CRISPR-based approach, the library incorporated sgRNAs targeting highly connected proteins (see Methods) defined by a stringent STRING protein-protein interaction (PPI) network (Figure 2A and Extended Data Table 1B)^28^ used to evaluate oncogenic signaling redundancies and interpret co-dependencies. Three out of four available PDX lines (PATC69, PATC124 and PATC53) were compatible with the CRISPR-based screening technology *in* vivo (Extended Data Figure 2B). We conducted each in vitro CRISPR screen as a time course, with a matched in vivo endpoint (Extended Data Figure 2A).

Data from the sgRNA screens were not amenable to analysis using current analytical frameworks (e.g. MaGeCK^29^, JACKS^30^, CERES^31^, and BAGEL^32,33^), which are designed for genome-wide screens and are not tailored for the small training sets of our custom library. To address this, we adapted the BAGEL framework^32,33^ to create Low-Fat BAGEL, which is optimized to analyze small targeted sgRNA libraries that are needed for *in vivo* screening or in other experimental settings where complexity may be a limitation. The difference in performance between BAGEL and Low-Fat BAGEL in dealing with outliers (Extended Data Figure 3A) is exemplified in the Bayes Factors (BF) (Extended Data Figure 3B and Extended Data Figure 3C) and Precision-Recall curves for a particular screen in PATC69 PDX lines (Extended Data Figure 3D). In comparison to other genome-wide analytical methods, Low-Fat BAGEL analyses demonstrate better performance and a more accurate classification of essential and nonessential genes in our screens (Extended Data Figure 3E). To ensure quality control, complexity coverage for RNAi and CRISPR libraries was confirmed for each in vitro and in vivo screen (Extended Data 4A - C). Control separation was confirmed in RNAi screens, and fold change separation and precision-recall of the 50 essential and 50 non-essential control populations were confirmed for each CRISPR screen using Low-Fat BAGEL (See Methods, Extended Data Figure 4D – I).

Quantile-normalized BFs (BF > 1) were directly compared to uncover essential genes^32^ represented in the three models *in vitro* and *in vivo* (Figure 2B and Extended Data Figure 5A). Dependencies among the three PDX lines, in both *in vitro* and *in vivo* conditions, were highly diverse. Ribosome-associated *RACK1*, *RPL30*, and the *MYC* proto-oncogene were the only shared vulnerabilities identified in all *in vivo* and *in vitro* screening contexts (Figure 2B and Extended Data Figure 5A). Quantile-normalized BFs also highlighted varying degrees of essentiality for *KRAS*, with PATC124 growth exhibiting less dependence on *KRAS* compared to PATC69 and PATC53 PDX lines (Figure 2C – E and Extended Data Figure 5B – D), which suggests that oncogenic RTK buffering may contribute to signaling complexity even in the context of mutated *KRAS*.

### CoDEX platform identifies EMT-associated dependencies along prominent PDAC molecular signatures

Next, we integrated quantile-normalized BFs with the STRING PPI network used to build the sgRNA library to generate dependency networks for each model. Dependency networks were merged across all three PDX lines based on overlapping essentiality thresholds (BF >1) for *in vivo* and *in vitro* contexts and displayed using a force-directed layout to visualize connectivity between established and novel gene targets (Figure 2C – E and Extended Data Figure 5B – D). The *in vivo* PATC53 dependency network highlighted multiple unique groups of interconnected vulnerabilities when compared to PATC69 and PATC124 dependency networks, despite PATC53 and PATC124 both being classified as basal-like, based on the Moffitt signature (Extended Data Figure 2B). Notably, interconnected vulnerabilities in PATC53 were associated with epithelial-to-mesenchymal transition (EMT) (e.g. *SNAI1*, *ZEB1*, *SMAD4*, *MAPK11* and *MAPK14*), integrin signaling (e.g. *ILK*, *ITGB1*, *ITGB2* and *NPNT*), heparan sulfate proteoglycan regulation (e.g. *SDC3* and *EXT1*), cell junction regulation (e.g. *CTTN*, *TJP1* and *TJP2*), and intracellular signaling kinases (e.g. *BCAR1*, *NCK1*, *CRKL* and *CRK*) (Figure 2F – H, Extended Data Figure 5E – G).

To anchor these vulnerability trends within the landscape of PDAC diversity, we then integrated the CRISPR screen-defined dependencies within our co-expression network. The integration displayed a clear shift in the PATC53-associated dependency spectrum along the Cluster 1 -to- Cluster 23 axis, with many of the dependencies identified in the CRISPR screen localized within, or adjacent to, Cluster 23 (Figure 1D, Figure 2F – H, Extended Data Figure 5E – G). Specifically, *MAPK11* and *FAM171A2* were localized within Cluster 23 itself; NKAIN4 in Cluster 25; and *SMAD4*, *ZEB1* and *SDC3* in Cluster 31 (Figure 1D, Figure 2F – H, Extended Data Figure 5E – G). The network localization of functionally annotated and PATC53-specific *ZEB1*, an EMT-associated transcription factor, and *SMAD4*, an EMT facilitator and oncogenic driver in advanced PDAC, implicated a connection between the Cluster 1-to-23 (C1vC23) axis, and its associated dependencies, with PDAC epithelial-to-mesenchymal transdifferentiation^34,35^.

### Prominent anticorrelated cluster signatures outline a continuous classical-to-basal differential signature

The distinct gene clusters we identified in the PDAC co-expression network provide a means to deeply characterize the diversity of any PDAC model by considering correlations in disease-specific cluster enrichment patterns, a parallel strategy to current approaches that use consensus clustering to assign a tumor subtype based on refined PDAC-specific gene sets. Thus, this refined co-expression network can comprehensively quantify global transcriptomic signaling trends on a tumor-by-tumor basis. Upon observing the clear dependency shift along Cluster 23 in the basal-like PATC53 model (Figure 2H), we noted a strong anti-correlative trend between Cluster 23 and Cluster 1 centroid scores among the entire PDAC PDX cohort (Figure 1E). The Cluster 1 centroid scores across our PDX cohort showed low variance among Moffitt-defined classical models, but a wider range and significant depletion of Cluster 1 gene expression (p = 3.67 × 10^−6^) was observed in basal-like models (Figure 3A). Interestingly, Cluster 1 unbiasedly localized the entire set of 21 classical signature genes from the Moffitt classification. Together, these findings suggested that we could leverage the anti-correlated C1vC23 axis as a classical to basal-like differential signature. Indeed, by applying K means clustering on only the Cluster 1 and Cluster 23 centroid scores, we separated the PDX cohort into three groups (Figure 3B): 1) enrichment in Cluster 1 represented Moffitt-defined classical models, 2) enrichment in Cluster 23 represented Moffitt-defined basal-like models, and 3) models with partial enrichment of both Clusters 1 and 23 (Figures 3C – D), which we termed “quasi-basal”. Thus, our C1vC23 signature provides a continuous transcriptomic signature that expands on the original binary Moffitt classification and uncovers a transitionary quasi-basal phenotype of PDAC with molecular signatures falling along a continuum.

**Figure 3:**
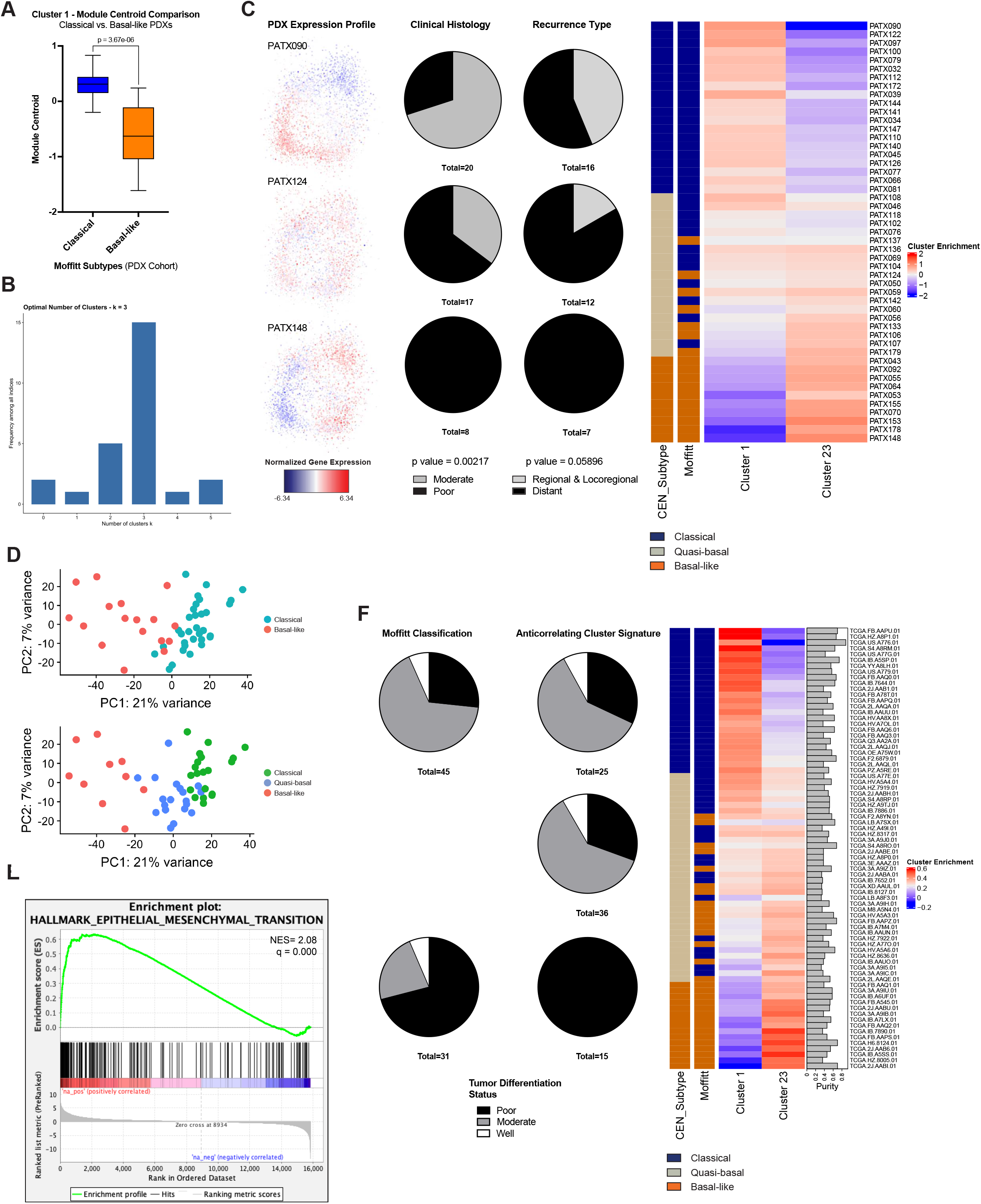
Prominent anti-correlated CoDEX clusters recapitulate a granular spectrum of “classical”, “quasi-basal”, and “basal-like” cells inter-tumorally. **(a)** [left panel] Overlay of the normalized expression score of each gene over the PDAC co-expression network of CoDEX defined classical, quasi-basal and basal models [middle panel]. Pie chart comparison of clinical histology and recurrence of patients associated with PDX models across subtypes. P values are derived from Chi Square test. [right panel] K means clustering reveals optimal k=3 using the Cluster 1 and Cluster 23 centroid scores across the PDX models paired with Consensus Clustering (k=2) into Moffitt’s classical and basal subtypes (**b**) Box plot of average Cluster 1 centroid score across Moffitt-defined classical and basal-like models. Box-whisker plots show median◻±◻first and third quartiles. P◻values are derived from t–test (n◻=◻48 PDX tumors). (**c**) PCA demonstrating the subtypes derived from the PDAC co-expression network with the quasi-basal group driving the PCs compared to the binary Moffitt Classification. (**d**) GSEA conducted on PDAC co-expression network-defined classical and basal-like models identifies EMT as a top pathway enriched in basal-like models FDR = 0.000. (**e**) [left panel] Pie chart comparison of tumor differentiation status in high-epithelial-content TCGA models comparing PDAC co-expression network-derived classifications and Moffitt classifications (n=76 tumors). [right panel] Heat map of C1vC23 anti-correlated cluster signature differential, matching Moffitt classification and tumor grade on TCGA tumor samples (samples with ≥ 30% epithelial content).

To gain further insight into the molecular signatures driving PDAC subtypes, we conducted gene-set enrichment analysis to compare CoDEX-defined classical vs. basal-like models within our PDX cohort. This highlighted that EMT signaling was significantly enriched among PDX models that fell into the CoDEX definition of basal-like, whereas this gene set was depleted in classical models (Figure 3E). Enrichment of an EMT signature in basal-like PDX models supports the hypothesis that the basal-like subtype is strongly associated with tumors where a majority of tumor cells has at least partly undergone transdifferentiation towards a mesenchymal phenotype. Consistently, analysis of clinical and histological data (see Methods) for the PDX cohort demonstrated that basal-like tumors were uniformly associated with poor differentiation status and distant metastatic recurrence (Figure 3B). Next, using high-epithelial-content PDAC patient data from TCGA (30% - 80% epithelial cells), we confirmed the presence of the network-derived subtyping, again identifying a quasi-basal continuum. Further, we recapitulated the association between tumor histology and the network-derived subtype in the TCGA dataset, wherein more poorly differentiated tumors were classified as basal-like, with strong enrichment in Cluster 23 gene expression (Figure 3F). By quantitatively characterizing a quasi-basal population, the CoDEX platform enables a more precise definition of the clinically relevant basal-like tumor cohort while also expanding on associated molecular dependencies that may be considered for therapeutic intervention^7^. Moreover, CoDEX defines a broader, cluster-level characterization of the range of molecular signaling contributing to diversity in PDAC, which can be used to granularly ascertain pathways that may be therapeutically targeted to exert anti-tumor effects across this classical, quasi-basal and basal-like tumor spectrum.

### Single-cell transcriptomic profiles of tumors define a cell-intrinsic clonal signature of PDAC subtypes

To investigate whether the quasi-basal signature identified in bulk tumor populations represents a quantifiable cell state or a mean signature derived from competing subcellular populations, we conducted single-cell RNAseq (scRNAseq) on the PATC69 (quasi-basal), PATC124 (quasi-basal), and PATC53 (basal-like) PDX lines. The co-expression network was used to identify cells expressing more than 30% of any cluster, and a centroid score for that cluster was calculated using genes with expression highly correlated to the cluster enrichment (r > 0.4). By analyzing scRNAseq data in the context of the co-expression network, we circumvented the technical issue of signal dropout by prioritizing large gene clusters to represent transcriptional diversity rather than single gene expression. Thus, the co-expression network serves as an additional resource for disease-specific single cell analysis. This approach successfully confirmed the presence of a quasi-basal C1vC23 signature as a quantifiable state in individual cells within in each PDX model (Figure 4A – D), confirming that the bulk readout represented an average of the intratumoral spectrum of classical, quasi-basal, and basal-like subclonal populations, rather than competing classical and basal-like signatures.

**Figure 4:**
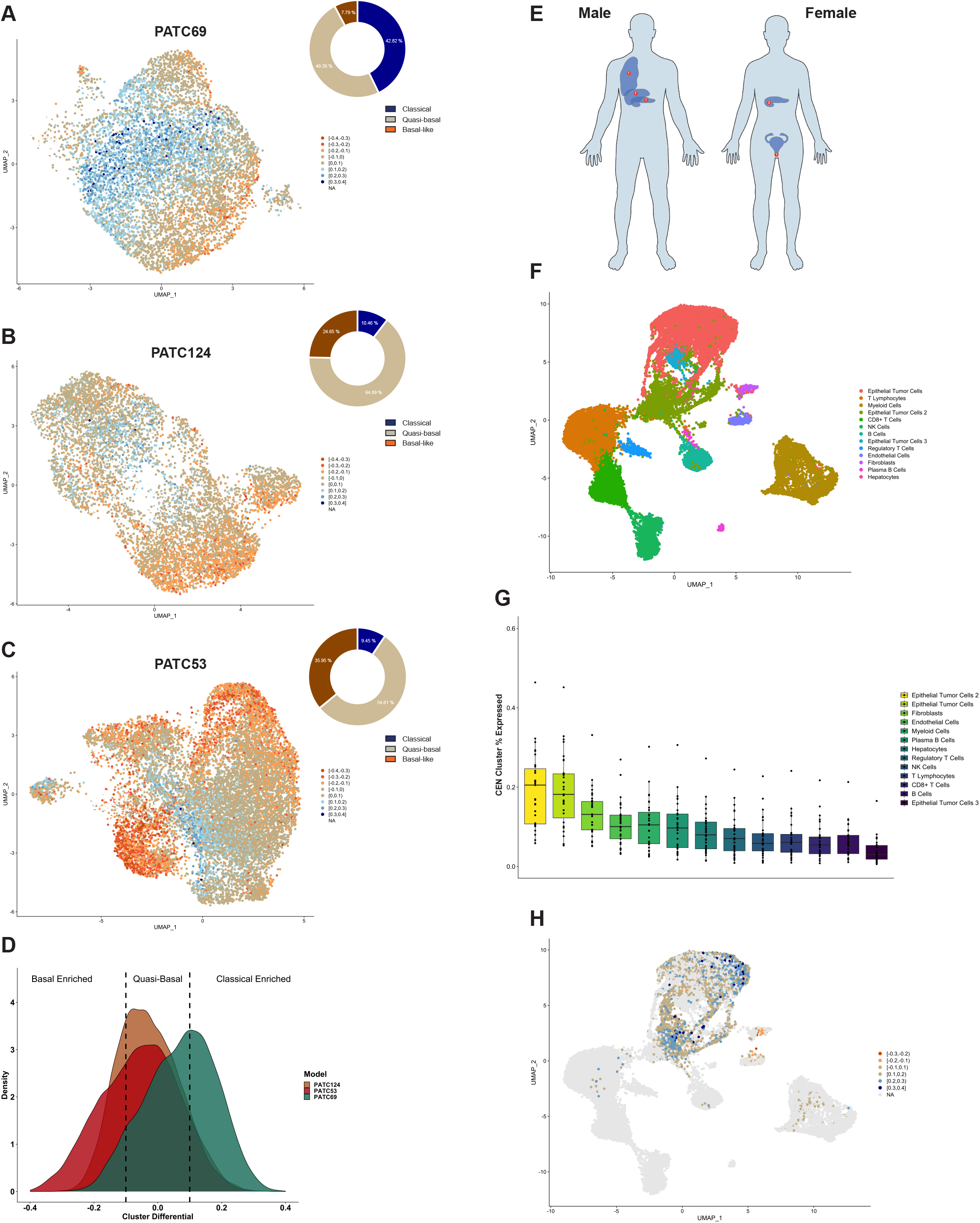
Intra-tumoral characterization of anti-correlated CoDEX clusters recapitulate a granular and tumor-localized “classical”, “quasi-basal”, and “basal-like” spectrum. (**a-c**) UMAP of single-cell sequenced low-passage (a) PATC69 (7,857 cells), (b) PATC124 (9,482 cells) and (c) PATC53 (14,791 cells) cell lines with an overlay of a PDAC co-expression network-normalized C1v23 signature differential with respective percentages of cells corresponding to each subtype [right panels]. Pie chart with distribution of three classifications (percentage) (**d**) Density histogram of the C1v23 signature differential distributions of PATC124, PATC53, and PATC69 PDX lines, with more positive cluster differential indicating enrichment in Cluster 1 and more negative cluster differential indicating Cluster 23 enrichment. (**e**) Diagram of the locations and numbers of isolated patient core needle biopsy (CNB) samples used for single-cell RNA sequencing analysis. (**g**) Combined UMAP outlining the tumor microenvironment components of multiple single-cell sequenced primary (n = 4) and metastatic (n = 3) CNB samples from PDAC patients. (**g**) The mean percentage of cluster representation for each cluster in the PDAC co-expression network from the cellular constituents of the tumor microenvironment. Each dot per cell type represents individual clusters. Box plot center: mean; box: quartiles 1–3; whiskers: quartiles 1−3 ± 1.5 × IQR. (**h**) Combined CNB UMAP with an overlay of a PDAC co-expression network-normalized C1vC23 signature represented as the difference between Cluster 1 and Cluster 23 expressions.

To evaluate the utility of the network-defined tumor subtypes in a clinical setting, in which small numbers of cells and stromal components may influence molecular analyses, we evaluated seven patient core needle biopsies (CNBs; four primary tumors, as well as one each of liver, lung, and vaginal metastases) using the network-aided scRNAseq analysis^36^ (Figure 4E, Extended Data Figure 6A). We first defined the distribution of cell types amongst our samples (Figure 4F) using previously annotated marker genes (Extended Data Figure 6B) and characterized cluster representation across the tumor microenvironment by quantifying the mean percentage of genes of each cluster for each cell type. This identified the epithelial component of the tumor as the primary contributor to the co-expression network cluster signatures (Figure 4G). Applying the same cluster centroid normalization method described above, the majority of cells that met our quality control cutoff were epithelial cells, with very little representation from two primary tumor samples (Primary 1 and Primary 2) that contained little to no epithelial content (Extended Data Figure 6A). Finally, to determine where each cell type was represented on the classical to basal-like continuum, we assessed the C1vC23 cluster differential. We found that the C1vC23 classification is largely present within epithelial cells, with Uniform Manifold Approximation and Projection (UMAP) clusters of fibroblasts and endothelial cells only representing a minority of misclassified C1vC23 signatures within the representative multiregional tumor microenvironment (Figure 4H). This finding confirms a lack of sample purity bias in the characterization of C1vC23 signatures in the TCGA PDAC samples (outlined in Figure 3F) and supports the feasibility of applying network-based cluster characterization to bulk clinical samples, including CNBs.

### CoDEX platform prioritizes subtype-associated biomarkers

Our PDAC-specific co-expression network allows us to leverage cluster centrality to associate cluster-representative biomarkers to signatures, and the CRISPR screens uncovered first-degree nodes of specific dependencies that can also serve as significantly co-expressed biomarkers. By jointly applying centrality and defining hit-specific nodes to prioritize biomarkers, we capture both broader, disease-specific cluster patterns as well as the expression of individual genetic vulnerabilities of potential interest. Accordingly, we used multiplex immunofluorescence (IF) to independently recapitulate the subtype-defining C1vC23 transcriptomic signature at the protein level. This method provides an avenue to quantitatively prioritize potentially useful target-associated biomarkers while also serving as a protein-level validation of this transcriptomic signature. As a proof of concept, we first determined closeness centrality for all genes in both Cluster 1 and Cluster 23 (Figure 5A). We then defined the first-degree nodes of two *in vivo* targets, *DNM2* and *MAPK11*, which localized to Cluster 1 and 23, respectively (Figure 5B – C). Among the first-degree nodes, we then prioritized genes in each gene set based on closeness centrality and antibody availability, which resulted in selection of *VSIG1* and *VIM* as representative biomarkers of Cluster 1 and 23, respectively (Figure 5D). IF analysis and image-based quantification of *VSIG1* and *VIM*, when combined with double staining for HLA, successfully recapitulated the expected C1vC23 signature at the protein level in PDX models with the representative classical (VSIG+), quasi-basal (VIM−/VSIG−), and basal-like (VIM+) transcriptomic signatures (Figure 5E, F). These findings demonstrate the utility of jointly leveraging cluster-centrality and defined gene targets to quantitatively prioritize protein biomarkers associated with relevant transcriptional profiles. Moreover, these biomarkers provide protein-level validation of transcriptomic cluster trends and serve as accessible alternatives for clinical characterization.

**Figure 5:**
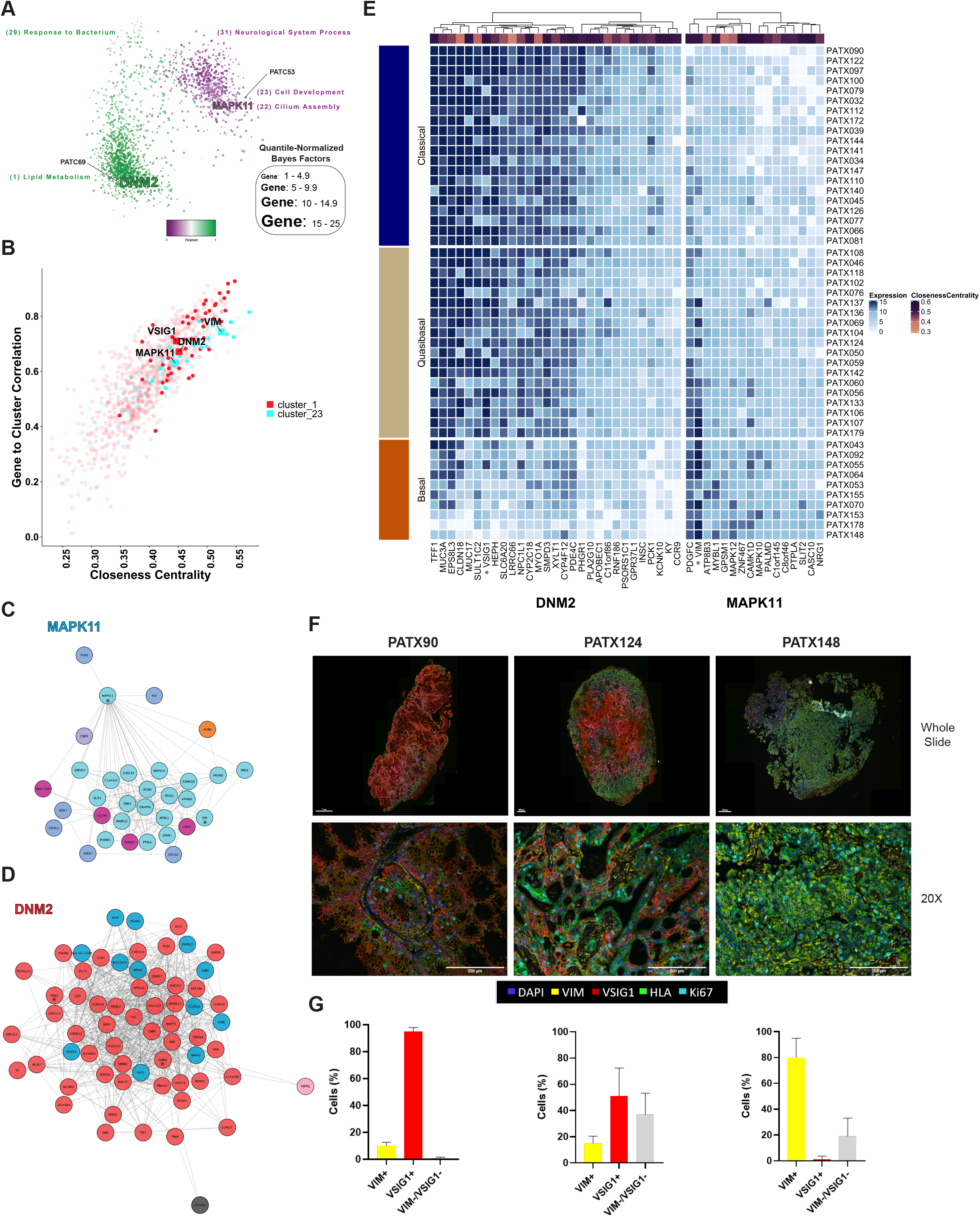
CoDEX platform prioritizes potential biomarkers by integrating first-degree nodes of anchored dependencies and cluster centrality. (**a**) Force-directed layout of prominent anti-correlative cluster centroid trends across the PDAC co-expression network highlighting Cluster 1 (Lipid metabolism) vs. Cluster 23 (Cell development) along with PATC69 *in vivo* specific vulnerability, DNM2, and PATC53 specific *in vivo* vulnerability, MAPK11, from our CRISPR screen analysis. Force-directed network outlining first-degree nodes of DNM2. (**b**) Dot plot of closeness centrality of each gene within Cluster 1 and Cluster 23 calculated using Cytoscape compared with gene-to-cluster correlation calculated as the Pearson correlation between single-gene expression and Cluster 1 and 23 centroids using the PDX cohort (n=48) (**c**) Force-directed layout of first-degree nodes of Cluster 23 localized MAPK11 generated from Cytoscape, or direct gene-wise correlations within the co-expression network with color of each gene representing its respective cluster assignment **(d)** Force-directed layout of first-degree nodes of Cluster 1 localized DNM2 generated from Cytoscape **(e)**Heat map of the normalized expression the first-degree node of DNM2 and MAPK11 in the PDX models (n=48), displayed along the C1vC23 signature-determined classification on the left and gene-wise closeness centrality on the top annotation (**f**) IHC-IF for VSIG1 vs. VIM, in the context of HLA, DAPI and Ki67, in representative slices of classical (PATX90), quasi-basal (PATX124) and basal-like (PATX53) PDXs (**g**) Quantification of VIM-positive, VSIG1-positive, and double-negative signatures in HLA-positive tumor cells.

### CoDEX-informed genetic dependencies functionally validate intratumoral classical, quasi-basal, and basal-like molecular signatures

By integrating CRISPR screen-defined co-dependencies with the co-expression network, the CoDEX platform can identify both common and unique dependencies within the larger context of PDAC diversity. To determine the functional relevance of this approach, we evaluated the effect of perturbing gene targets associated with the basal-like subtype that were prioritized through our CoDEX analysis in varied tumor contexts (Figures 2F – H, Figure 6A). Based on our *in vitro* CRISPR screening results, we selected sgRNA sequences targeting *SMAD4*, *ZEB1*, and *ILK*, as well as the non-essential gene, *ABCG8, as* a negative control (Figure 6B, Extended Data Figure 6A). CoDEX-informed *SMAD4* and *ZEB1*, both localized within the network, were targeted for C1vC23 signature validation. *ILK*, a CRISPR-defined dependency in PATC53 not present in the co-expression network, was selected to determine whether knockout of this potential EMT regulator would also have the capacity to influence the C1vC23 signature differential^37–39^. In addition, we applied a feature barcoding strategy whereby a complement sequence was incorporated into the 3’ end of the sgRNA sequences, enabling scRNAseq sample multiplexing and quantification of the C1vC23 signature shift relative to the sg*ABCG8* negative control distribution (Figure 6B). Individual sgRNAs derived from the CRISPR library were transduced into quasi-basal PATC69 and basal-like PATC53 cells (Extended Figure 7A). Cells were cultured cells *in vitro* and collected at the earliest point when separation of essential versus non-essential genes was observed in the original CRISPR screen (Day 20) (Figure 6B and Extended Data Figure 4D – H). Sanger sequencing was used to analyze the indel frequency of each sgRNA (Extended Data Figure 7B - K), and colony growth was tracked for each sgRNA to confirm selective growth inhibition in the basal-like PATC53. (Figure 6B). Multiplexed scRNAseq was conducted on 10,000 cells total (2,500 cells per sgRNA) for each PDX line model.

**Figure 6.**
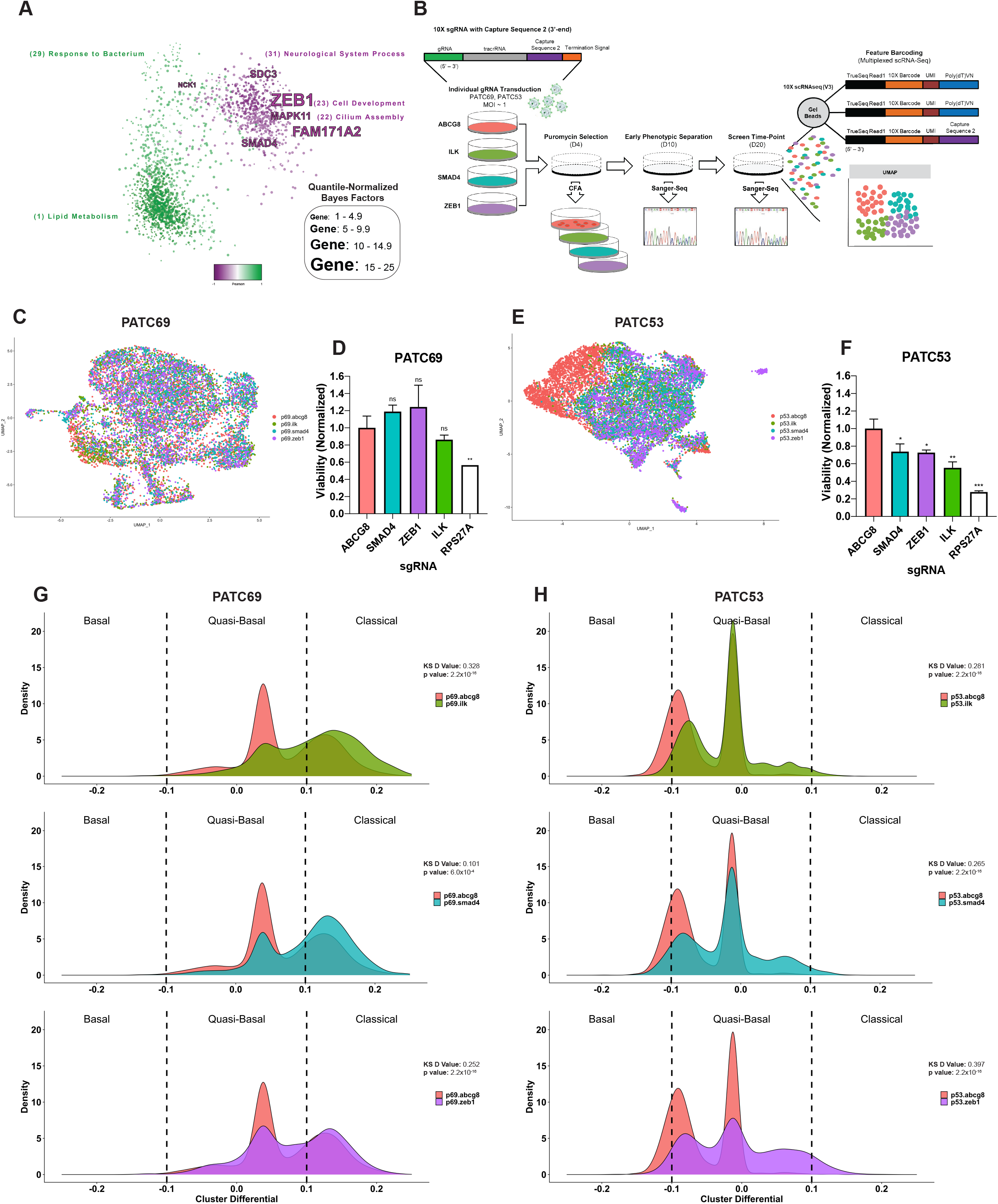
Functional validation of CoDEX-informed intra-tumoral context dependency for C1vC23-associated genetic targets. (**a**) Force-directed layout of prominent anti-correlative cluster centroid trends across the PDAC co-expression network highlighting Cluster 1 (Lipid metabolism) vs. Cluster 23 (Cell development) along with PATC53 *in vivo* specific vulnerabilities from our CRISPR screen analysis with size of the gene representing the quantile-normalized BF (**b**) Overview of the feature barcoding strategy for intratumoral tracking of PDAC co-expression network signatures following sgRNA knockouts. Targeting sgRNAs for *ABCG8* (as a negative control), *ILK*, *SMAD4*, and *ZEB1* were selected and based on CRIPSR screening results. PATC69 and PATC53 cell populations were individually transduced *in vitro*, selected with puromycin, and parallel assays were also prepared to confirm knockout phenotypes through colony growth, and sgRNA cutting through Sanger sequencing. Each knockout cell population was cultured *in vitro* for 22 days (CRISPR screening time-point day 10), and then combined scRNAseq library preparation. A total of 10,000 cells per PDX line were sequenced, 2,500 per condition. (**c**) UMAP of the PATC69 PDX line with defined *ABCG8*, *ILK*, *SMAD4*, and *ZEB1* knockout populations (10113 total cells). (**d**) Normalized viability of PATC69 cells following knockout of *ABCG8*, *SMAD4*, *ZEB1*, *ILK*, or *RPS27A* with sgRNA. **p < 0.05. (**e**) UMAP of the PATC53 PDX line with defined *ABCG8*, *ILK*, *SMAD4*, and *ZEB1* knockout populations (11,439 total cells). (**f**) Normalized viability of PATC53 cells following knockout of *ABCG8*, *SMAD4*, *ZEB1*, *ILK*, or *RPS27A* with sgRNA. *p < 0.05; **p < 0.01. (**g**) Comparative PATC69 density plots of C1vC23 signature following knockout with sg*ILK*, sg*SMAD4*, sg*ZEB1* and sg*ABCG8* negative control calculated as the differential between the Cluster 1 and 23 centroid score, with more positive differential indicating cluster 1 enrichment and vice versa. P values and D statistic derived from Kolmogorov Smirnov test (**h**) Comparative PATC53 density plots of C1vC23 signature in PATC53 cells following knockout of *ILK*, *SMAD4*, *ZEB1*, or *ABCG8* (negative control).

PATC53 UMAPs revealed a clear separation between the *ABCG8* negative control knockout cells and populations with perturbations in combined Cluster 23 (*SMAD4* and *ZEB1*) and basal-like (*ILK*)-associated genes, whereas no separation was observed between test genes and the negative control in PATC69 cells (Figure 6C, E). Also, as expected, the PATC69 population transduced with negative control sg*ABCG8* contained C1vC23-defined classical and quasi-basal cells, while the basal-like PATC53 population contained quasi-basal and basal-like cells (Extended Data Figure 8C - D). To test the C1vC23 signature distribution shift relative to the reference distribution defined by the *ABCG8-*null negative control, C1vC23 density plots were generated for each sgRNA knockout cell line (Figure 6G – H), and Kolmogorov’s D statistic was used. Deletion of *ILK*, *SMAD4*, and *ZEB1* each resulted in a significant C1vC23 shift towards the Cluster 1-enriched classical signature in both PATC69 and PATC53 (Figure 6G – H). In parallel with scRNAseq, transduced populations were seeded *in vitro* immediately following selection to compare relative growth phenotypes for both PDX lines (Figure 6B). Coinciding with the signature shift towards Cluster 1 at the single-cell level, perturbation of each of the three basal-like-associated genes inhibited bulk population growth in PATC53 relative to sg*ABCG8*-knockout controls, whereas no growth phenotype was observed in the classical PATC69 (Figure 6D, F and Extended Data Figure 8A, B).

In summary, we provide functional validation of the C1vC23 signature based on concordant signature shifting, in both quasi-basal PATC69 and basal-like PATC53 cell populations, following genetic perturbation of basal-like associated targets. Importantly, these findings provide evidence for our CoDEX platform to uncover the intratumoral context of vulnerabilities localized within and adjacent to Cluster 23, by quantifying the targeted depletion of C1vC23-defined basal-like cell populations. This represents a novel approach to leverage network-informed signaling perturbation to influence heterogeneity in pancreatic tumors.

## DISCUSSION

PDAC characterization and subtyping centered on the categorical Moffitt classical and basal-like molecular subtypes are able to stratify patients based on clinical outcome, but have not yet been able to guide precision medicine^7^. We developed CoDEX as an unbiased platform to quantify the molecular signatures that contribute to PDAC diversity and provide signaling context to functionally annotated genetic dependencies. By applying the CoDEX platform to our cohort of PDAC PDX models and patient samples, we identified a differential signature derived from anti-correlated Clusters 1 and 23, C1vC23, which demonstrated a classical-basal continuum corresponding with EMT. GO annotations underlying the identified clusters provided novel insight into the oncogenic signaling characteristics of the Moffitt-defined classical and basal-like subtypes. Cluster 23, which defined basal-like subtypes, was characterized by gene signatures associated with cell projection organization, cell motility, and general cell development. Alternatively, Cluster 1, which defined classical tumors, exhibited a transcriptomic bias towards lipid metabolism, hormone regulation, and other xenobiotic metabolism signaling genes. Lipid metabolism and lipid droplets have previously been observed to play important roles in generating the fatty acids required for pancreatic cancer cell proliferation^40^. Thus, linking *de novo* lipid synthesis to the classical tumor spectrum provides key insight into potential context-specific metabolic dependencies in lower histological grade PDAC^40^.

Application of the C1vC23 signature yielded a notable refinement in the Cluster 23-enriched basal-like cohort that was defined by a significant enrichment in mesenchymal-associated clinical features and genetic dependencies. This basal-like refinement was due to the quantification of a near-zero differential in the C1vC23 signature at the bulk tumor and single-cell levels, which we defined as quasi-basal. Importantly, this quasi-basal single-cell population was significantly represented within both bulk classical and basal-like tumors, expanding on the Toronto classification approach that demonstrated co-occurrence of classical and basal-like subtype heterogeneity at the single-cell level^10,41^.

In addition to quantifying the transcriptomic diversity within PDAC, the CoDEX platform integrates prioritized *in vivo* functional genomics on our PDX samples that results in a clinically relevant, context-forward approach to contextualize the nature of genetic dependencies. By using PDXs, we overcome the lack of Moffit-defined classical cell lines available for PDAC research^7^. scRNAseq analysis of phenotypically and genetically diverse PDX models recapitulated findings from bulk tumors, revealing vast intratumoral heterogeneity, and a cluster-defined classical to basal-like clonal spectrum with a discrete quasi-basal signature in individual cells. Furthermore, scRNAseq of patient samples recapitulated the C1vC23 intratumoral spectrum and revealed that the majority of the co-expression network signatures was intrinsic to tumor cells.

We validated our C1vC23 signature by directly quantifying the effect of informed target perturbation on C1vC23 subclonal heterogeneity through feature-barcoded scRNAseq. Multiplexed CRISPR deletion of basal-like associated and cluster 23-anchored targets, *ILK*, *SMAD4*, and *ZEB1* in knockout cells quantified a “subclonal shift” toward the classical signature. This change in molecular dominance, observed in both basal-like and quasi-basal populations, functionally demonstrated context-specific gene essentiality as a function of disease evolution. Furthermore, knockout of established mesenchymal drivers, such as *SMAD4* and *ZEB1*, served to validate the C1vC23 signature. Consistent with previous studies, our findings support a dual function of SMAD4 in PDAC, and the CoDEX-defined C1vC23 signature puts the tumor suppressor and tumor promotor roles of SMAD4 within distinct molecular contexts^35,42^.

Our integrated CoDEX approach introduces multiple avenues for hypothesis generation within the context of tumor heterogeneity. For example, anchoring of relatively less studied targets (*e.g.*, *FAM171A2* and *NKAIN4*) to Cluster 23 implicates context-specific roles in mesenchymal cell proliferation and maintenance. Although our results here focused on selective gene dependencies within highly aggressive basal-like sub-clones, functional maps developed through the CoDEX pipeline also unraveled Cluster 1-associated vulnerabilities, such as *DNM2*, with potential to eradicate classical clonal subpopulations. Thus, while the quasi-basal population exhibits a near-zero differential in the C1vC23 signature, associating other network-derived cluster signatures (e.g. C2vC13) may be useful to identify selective genetic dependencies and further stratify this heterogeneous sub-population. This context-forward approach to stratify dependencies provides a novel avenue to expand on current PDAC subtypes and to define gene targets localized within additional cluster patterns underlying PDAC heterogeneity.

Notably, the subclonal shift induced by *ILK* knockout highlights the fact that additional metrics at the protein-level are needed to more comprehensively quantify PDAC diversity, and underscores the need to integrate protein status with network cluster trends to understand how molecular signaling is altered or repositioned following major transcriptomic rewiring. Future integration of the co-expression network with partially recapitulated or intact immune and stromal components may also elevate functional characterization from the cell-intrinsic to tumor-intrinsic setting.

The use of CoDEX as an integrated approach to deconvolute tumor heterogeneity into relatively more homogenous and approachable cluster-defined subclonal populations may, ideally, help to direct treatment options to more significantly and comprehensively impact pancreatic tumors. This strategy may also inform on the implementation of novel or existing drugs that may mitigate tumor heterogeneity, but have only a negligible effect on tumor volume and viability. To our knowledge, CoDEX serves as the first platform to define and leverage a comprehensive set of transcriptional signatures as a foundation to inform context-specific dependency.

## METHODS

### PDX models and Sequencing (RNA and Whole Exome)

A total of 49 models were utilized in this paper. PDAC PDX models were obtained from the labs of Dr. Michael Kim (Department of Surgical Oncology, MD Anderson Cancer Center) and Dr. Scott Lowe (Memorial Sloan Kettering Cancer Center)^43,44^. PDXs were propagated and maintained in NOD scid gamma (NSG) mice carrying NOD.Cg-*Prkdc*^*scid*^ *Il2rg*^*tm1Wjl*^/SzJ (Jackson Labs).

### PDX Sequencing

#### Whole exome library preparation and sequencing

Whole exome sequencing (WES) libraries were prepared using the Agilent SureSelect XT library preparation kit in accordance with the manufacturer’s instructions. Briefly, DNA was sheared using a Covaris LE220. DNA fragments were end-repaired, adenylated, ligated to Illumina sequencing adapters, and amplified by PCR. Exome capture was performed using the Agilent SureSelect XT v4 51Mb capture probe set and captured exome libraries were enriched by PCR. Final libraries were quantified using the KAPA Library Quantification Kit (KAPA Biosystems), Qubit Fluorometer (Life Technologies) and Agilent 2100 BioAnalyzer and were sequenced on an Illumina HiSeq2500 sequencer using 2 × 125bp cycles. Base calling and filtering were performed using current Illumina software and adapters were trimmed using Trim Galore [55]. Sequences were aligned to both NCBI genome human build 37 and mouse build 38 using Burrows-Wheeler Aligner^45^; identified mouse reads were removed from the original FASTQs and then the files were realigned again to NCBI build 37 using BWA. Picard was used to remove duplicate reads (http://picard.sourceforge.net); base quality scores were recalibrated using GATK^46^. Assessment of reads that do not align fully to the reference genome was performed, locally realigning around indels to identify putative insertions or deletions in the region. Variants were called using GATK HaplotypeCaller, which generates a single-sample GVCF. To improve variant call accuracy, multiple single-sample GVCF files were jointly genotyped using GATK GenotypeGVCFs, which generates a multi-sample VCF. Variant Quality Score Recalibration (VQSR) was performed on the multi-sample VCF, which adds quality metrics to each variant that can be used in downstream variant filtering.

#### RNA library preparation and sequencing

RNA sequencing libraries were prepared using the Illumina TruSeq Stranded mRNA sample preparation kit in accordance with the manufacturer’s instructions. Briefly, 100ng of total RNA was used for purification and fragmentation of mRNA. Following conversion of mRNA to cDNA, DNA was adenylated, ligated to Illumina sequencing adapters, and amplified by PCR (using 10 cycles). Final libraries were quantified using the KAPA Library Quantification Kit (KAPA Biosystems), Qubit Fluorometer (Life Technologies) and Agilent 2100 BioAnalyzer and were sequenced on an Illumina HiSeq2500 sequencer (v4 chemistry) using 2 × 50bp cycles. Base calling and filtering were performed using current Illumina software. Reads were aligned to a joint index of NCBI genome human build 37 and mouse build 38 with STAR aligner^47^. Reads that map uniquely and unambiguously to the mouse genome were removed from the FASTQ files and then the files (containing unmapped reads and reads mapped at least once to the human reference) were remapped to GRCh37 using STAR aligner and Gencode 19 annotation. Gene expression quantification was performed with featureCounts (http://bioinf.wehi.edu.au/featureCounts/). Genes with raw read counts present in less than 20% of samples were removed from further analysis, and log normalized counts were generated on the 14175 filtered genes using DESeq^48^.

### Moffitt Classification

Moffitt classification of Classical and Basal was assigned to the 49 PDX models using log normalized and scaled counts of 21 classical and 25 basal genes for PDX PDAC model classification described by consensus clustering into two groups using the R package ConsensusClusterPlus^49^. The clusters were manually assigned as classical and basal based on high expression of each group of genes.

#### Construction of the PDAC co-expression network

Gene-wise median absolute deviation across samples from the normalized read counts. 8505 genes with median absolute deviation exceeding the 40^th^ percentile was filtered to consider highly variable genes. A comprehensive network of all 8505 was generated by calculating spearman correlation across all gene pairs. In order to prune the network to prioritize biologically relevant gene pairs, a Bayesian framework developed by Yang, et al, Log Likelihood Score (lls), was applied^15^. This paper curated a “positive gold standard” list of biologically relevant gene pairs and “negative gold standard” gene pairs with no known functional annotations. The likelihood of correlation between any given gene-pair being functionally relevant is calculated comparing to the negative and positive gold standard gene pairs. A log likelihood score of 2.5 was used to cut off gene pairs to be included in the network.

#### Clustering and GO pathway annotation

Infomap^16^, a community detection tool for large networks, was used to cluster the network. All gene pairs were inputted into Infomap, and three hierarchical tiers of clusters were produced. In order to assign clusters, if the third tier contained more than 50 genes, it was assigned to an individual cluster. In the event that the third tier contained less than 50 genes, it was folded into the second tier, all of which formed a cluster. This was the case with cluster 29-31, which are larger encompassing clusters. For the resulting 31 clusters, the R package goseq^50^ was used to conduct hyper-enrichment analysis of Gene Ontology Biological Processes pathways on each cluster. The R package “revigo”^51^ was used to prioritize and visualize GO pathways to represent their hierarchical class.

### Mutation analysis

The “oncoplot” feature within the R package maftools^52^ was utilized to visualize the mutational spectrum across the genes identified as relevantly mutated by the PDAC TCGA paper. To identify complementary patterns of mutations across clusters, we used a method called UNCOVER^17^ using a filtered set of “high impact” mutations and the cluster enrichment centroid scores per model. The filtered set of mutations was generated by identifying canonical mutations with gnomAD^53^ allele frequency less than 1%, “moderate” and “high” impact, and limited to non-intronic/non-coding and synonymous mutations.

### Centroid scores

To generate a cluster level enrichment score, we calculated a centroid score per cluster by taking the mean log-normalized expression of all genes in each cluster for each sample. A comprehensive overview of the 31 centroid scores across the PDX models was generated using the R package ComplexHeatmap^49^. We further computed a Pearson correlation across all 31 clusters and identified strongly anticorrelated signature between Clusters 1 and 23.

### Network Display

Cytoscape (version 3.7.2) was leveraged for visualization of both the PDAC co-expression network and STRING-anchored co-dependency networks^54,55^. The PDAC co-expression network was visualized using a Prefuse Force-Directed Layout, with node color displayed based on Random Walk defined clusters and node size representative of Betweenness Centrality. For visualization purposes, only nodes with 12 or more edges are represented, and edges are not displayed in representative co-expression network figures. For STRING-anchored co-dependency networks, a Perfuse Force-Directed Layout is also applied, with quantile-normalized Bayes Factors represented as a blue -to- red color distribution. For each *in vivo* and *in vitro* condition, networks were constructed for each independent PDX line and then merged for comparison based on overlapping nodes. All STRING-defined edges were maintained for *in vivo* and *in vitro* network merging.

### Cell Culture

PDX cell lines were seeded in treated tissue-culture plates (Corning) in DMEM/F12 medium (Gibco) supplemented with 10% Fetal Bovine Serum (Gibco), Penicillin (50 units/mL) and Streptomycin (50 μg/mL) (ThermoFischer Scientific). Phosphate Buffered Saline (PBS) was utilized prior to trypsinization and for general cell washing purposes (Thermo Fisher Scientific). Cells were regularly trypsonized (0.25%, Trypsin-EDTA, Gibco) prior to reaching 70% - 80% confluence, and maintained on 10 cm and 15 cm treated tissue-culture dishes (Corning). Viable cells were counted using a Cellometer mini and 0.2% Trypan Blue staining (Nexcelom).

#### Design and Construction of Custom RNAi Library

The original RNAi custom library, constituted by 2,653 shRNAs targeting PDAC-prioritized proteins associated to the extracellular face of the plasma membrane, was constructed using chip-based oligonucleotide synthesis and cloned into a pRSI17cb-U6-sh-13kCB18-HTS6-UbiC-TagGFP2-2A-Puro lentiviral vector (Cellecta) as a pool. PDAC-prioritized surface proteins (including proteins with evidence of mislocalization) were selected based on tumor-specific overexpression compared to matched-normal tissue, copy number vs. RNA expression correlation, and SILAC screening for KRAS dependency^26^. Significant tumor-specific overexpression and copy number vs RNA expression were determined for the full transcriptome, and subsequently refined using an internally curated list of select extracellular proteins containing transmembrane domains, GPI-anchored proteins and proteins with evidence of membrane mislocalization. Tumor-specific overexpression vs. matched normal tissue was defined based on two matched-normal bulk tumor microarray datasets (FC ≥ 1.5, q < 0.005, ArrayExpress: E-GEOD-15471, EG-GEOD-28735), and further refined for tumor-specific expression (FC ≥ 1.5) by using a microarray dataset composed of micro-dissected samples (ArrayExpress: E-MEXP-1121/950)^23–25,56,57^. Positive Spearman correlations (rho ≥ 0.35, q < 0.005) of copy number vs. RNA expression were calculated based on available TCGA level 3 data derived from the 07/15/2014 dataset^58^. False discovery rate for bulk tumor microarray datasets and for copy number vs. RNA expression correlations was calculated using the “qvalue” Bioconductor package^59^. Genes derived from published data characterizing KRAS dependent surface protein localization, conducted across three iKRAS p53^L/+^ mouse^60^ tumor-derived cell lines, were also incorporated into the library based on previous work^26^. The shRNAs targeted 241 genes, with 10 shRNAs/gene. Targeting sequences were designed using a proprietary algorithm (Cellecta). The oligo corresponding to each shRNA was synthesized with a unique molecular barcode (18 nucleotides) for measuring representation by NGS. Negative controls consisted of shRNAs targeting Luciferase, and positive controls consisted of shRNAs targeting *RPL30* and *PSMA1*. In addition, we incorporated shRNAs targeting *KRAS* into the library to serve as additional controls for the PDAC PDX lines.

#### RNAi Screening *in vivo* and *in vitro*

Using the custom barcodes lentiviral shRNA library, PDX lines (PATC69, PATC124, PATC53 and PATC153) were transduced *in vitro* using 8 μg/mL Polybrene (Sigma-Aldrich). Libraries were transduced at 1000X coverage and multiplicity of infection (MOI) of 0.3. Media was replaced after 14 hours, and at 48 hours, and MOI was confirmed by checking GFP percentage with flow-cytometry 48 hours post-transduction. Immediately following MOI confirmation with flow cytometry, 2 μg/mL Puromycin (Thermo Fisher) was added to each transduced cell population for 72 hours. Immediately following Puromycin selection, a Reference population was collected (1000X) and stored at −80 °C. Remaining cells were split and moved into *in vitro* and *in vivo* settings. Three independent *in vitro* screens were seeded (1000X) into 15 cm treated plates (Corning). For *in vivo* implantation, cells were combined into 1:1 PBS/Growth-Factor Reduced Matrigel (Corning), and injected orthotopically at 1000X coverage per mouse. Immunodeficient NOD SCID mice were leveraged for *in vivo* screening. For *in vitro* screening populations, cell populations were collected at 10, 20 and 30 days post Reference collection. For *in vivo* screening, the entire pancreas and tumor of each mouse was collected at day 30. For each condition, cells were lysed using SDS and DNA was sheared using sterile 23 gauge 1 inch needles (Becton Dickinson). Tumor DNA was isolated using Phenol:Chloroform extraction and Ethanol precipitation. A Nested PCR strategy was utilized to amplify and prepare barcode populations for next-generation sequencing (Cellecta). Redundant siRNA activity (RSA) analysis was applied to rank targets for each PDX model and screen setting^61^.

#### Design and construction of custom CRISPR Library

The custom CRISPR-Cas9 sgRNA library was constituted by 3,367 sgRNAs and designed to both incorporate and expand upon the original RNAi-defined targets. The library was constructed using chip-based oligonucleotide synthesis and cloned into a pRSG16-U6-sg-HTS6C-UbiC-TagRFP-2A-Puro lentiviral vector (Cellecta) as a pool. RSA was used to rank essential genes from shRNA screens. Genes were curated based on RSA < 0.05 and FC < −2 in at least one model (N=100). Curated gene targets were further annotated by incorporating neighbors with a PPI score ≥ 0.80 (STRING, version 10), and TPM (transcripts per million) > 2 in that model^47^. In addition, 50 non-essential genes and 50 essential genes were added to have a final set of 654 genes^32^.

#### CRISPR Cas9 Screening *in vivo and in vitro*

Using the custom barcodes lentiviral sgRNA library, PDX lines (PATC69, PATC124, PATC53 and PATC153) containing the lentiCas9-blast vector (addgene, plasmid #52962) were transduced *in vitro* using 8 μg/mL Polybrene (Sigma-Aldrich). PDX lines transduced with Cas9 were constantly kept at 10 μg/mL blasticidin (Thermo Fischer Scientific). Libraries were transduced at 1000X coverage and a multiplicity of infection (MOI) of 0.3. Media was replaced after 14 hours, and at 48 hours, and MOI was confirmed by checking RFP percentage with flow-cytometry 48 hours post-transduction. Immediately following flow cytometry, 2 ug/mL puromycin (Thermo Fisher) was added to each transduced cell population for 72 hours. Immediately following puromycin selection, a Reference population was collected (1000X) and stored at −80 °C. Cells were allowed to culture for an additional 12 days in order to allow for some initial sgRNA-directed cutting prior to starting the *in vivo* screen, to reduce potential noise derived from disproportional cell doubling following orthotopic implantation. A secondary Reference (1000X), at the time of injection, was then collected. Remaining cells were split and moved into *in vitro* and *in vivo* settings. Three independent *in vitro* screens were seeded (1000X) into 15 cm treated plates (Corning). For *in vivo* implantation, cells were combined into 1:1 PBS/Growth-Factor Reduced Matrigel (Corning), and injected orthotopically at 1000X coverage per mouse. Immunodeficient NSG mice were leveraged for *in vivo* screening. For *in vitro* screening populations, cell populations were collected at 10, 20 and 30 days post injection and secondary Reference collection. For *in vivo* screening, the entire pancreas and tumor of each mouse was collected at day 30 post injection. DNA extraction and barcode library preparation were conducted in similar fashion to the RNAi screens. For each condition, cells were lysed using SDS and DNA was sheared using sterile 23 gauge 1 inch needles (Becton Dickinson). DNA was isolated using Phenol:Chloroform extraction and Ethanol precipitation. Nested PCR was utilized to amplify and prepare barcode populations for NGS (Cellecta).

### Low Fat BAGEL

Log2 fold-change was calculated on a guide level by comparing each time point to the reference time point for each model. Screen quality and efficient drop out was assessed by comparing the log density ratio log2 fold-change of core essential versus non-essential guides.

Low-Fat BAGEL is an adapted framework of the BAGELv2^33^ algorithm to more accurately analyze small screens with limited training sets. The BAGEL and BAGELv2 algorithms are previously described implementations of a Bayesian model selection algorithm, and in short, calculate a “Bayes Factor”, a log likelihood of gene essentiality trained on gold-standard core essential and non-essential genes^32^.

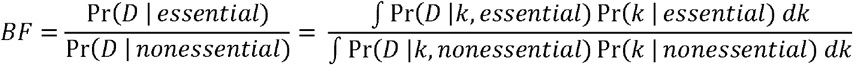

In Low-Fat BAGEL, a random bootstrap of core essential and non-essential guides is used as the training set to generate log density curves of the essential and non-essential fold-change distribution. This distribution is used to generate a linear regression model (k) of the fold-change to control for the influence of outliers. Over 100 permutations of the training set, Low-Fat BAGEL calculates the BF of all guides in the library, by comparing the log2 fold-change of each guide (D) to the distribution of fold-changes of the essential and non-essential genes. The final BF each of individual guide is computed as the mean across 100 iterations, and a gene level BF was calculated as the sum of all guides for that gene.

The R package ‘ROCR’ was used to generate Precision-Recall curves of the essential and non-essential gene level BF^62^. The F measure of each screen using different analytical methods was calculated as described here^32^.

#### Anticorrelated cluster signatures and “Quasi-basal” group

A test was performed to quantify the difference in Cluster 1 enrichment across Classical and Basal models. The centroid scores for Cluster 1 and 23 were used to categorize the PDX models into three clusters using K-means clustering. K=3 was chosen as the optimal number of clusters based on the majority consensus of two out of three methods between the elbow criterion in within-cluster sum of squares, BIC calculated by the R package “mclust”^63^ and “NbClust”^64^ which calculates the optimal k across 30 methods. The R package “factoextra”^65^ was used to visualize the results. The group of models with Cluster 1 enrichment and Cluster 23 depletion were termed “classical”, with the inverse group termed “basal”, and the third group with roughly equal centroid scores in Cluster 1 and 23 termed “quasi-basal”.

#### Differential Expression between Classical and Basal Models

After categorizing models, the ones deemed classical and basal using the CEN methodology were used to conduct differential expression analysis. DESeq^48^ was used to identify significantly differentially expressed genes, and genes were ranked by t score. GSEA was conducted using the “GSEA pre-ranked” feature^66^. We compared differences in histology and site of recurrence obtained from clinical data associated with each PDX model. Chi square tests were conducted using the CEN subtypes and Moffitt subtypes to evaluate if subtype level differences were statistically significant.

#### Clinical correlations with signature

Clinical data from patients corresponding to the PDAC PDX cohort was provided by the lab of Dr. Eugene Koay^67^. Differentiation status on 45 models was categorized as “Poor” and “Moderate” and histology status on 35 models was categorized as “Locoregional” if the records indicated as “regional” or “locoregional”, and “Distant” if site was outside the pancreas. Chi square tests were used to evaluate differences between the cluster-defined Classical, Quasi-basal and Basal groups as well as the Moffitt binary classification.

### TCGA data

TCGA gene expression data and corresponding clinical data was downloaded from GDC, and processed as previously described^4^. Raw read counts were log2 transformed and normalized using DESeq^48^ similarly to the PDXs described above. We filtered the dataset to “high purity” samples annotated in the clinical data, where we obtained the Moffitt classification data. Cluster centroids were calculated as described above with the PDXs and a k means of 3 was applied to the Cluster 1 and 23 centroid scores to categorize tumors into Classical, Quasi-basal and Basal. Tumor grade and differentiation status were obtained from the clinical data and Chi square tests were conducted to assess subtype level associations with the C1vC23 signature and Moffitt groups.

#### Single Cell Data for PDX-derived Cell Lines

Seurat version 3.1^68^ was used to analyze all single cell analysis. Each of the lines, PATC124, PATC53 and PATC69 were analyzed separately. PATC53 contained two replicates, which were merged for analysis. For all PDX cell lines, single cells with a minimum of 350 expressed genes and less than 10% mitochondrial reads were retained. Genes expressed in less than 3 cells and mitochondrial genes were removed from further analysis. The data was log normalized, transformed using the “vst” function with top 2000 variant genes. The total RNA count, cell cycle score and mitochondrial reads were regressed out. For PATC124, principal-component analysis and uniform manifold approximation and projection (UMAP) with the first 15 dimensions was performed, followed by identifying clusters using a resolution of 0.15. For PATC69, principal-component analysis and uniform manifold approximation and projection (UMAP) with the first 15 dimensions was performed, followed by identifying clusters using a resolution of 0.10. PATC53 contained two replicates, which were integrated as one dataset following normalization and variant stabilization. Total RNA count, cell cycle score and mitochondrial reads were regressed out of the integrated dataset, and PCA and UMAP on first 15 dimensions was performed, further identifying clusters using a resolution of 0.10.

#### Core Needle Biopsy Single Cell Analysis

Seven CNB samples were used – four primary tumors, one liver, lung and vaginal metastases sample each^36^. All seven samples were filtered to have a minimum of 350 expressed genes and less than 25% mitochondrial reads were retained. Similar to the PDX analysis, genes expressed in less than 3 cells and mitochondrial genes were removed from further analysis. Each sample was log normalized, transformed using the “vst” function with top 2000 variant genes. The seven samples were then integrated, following which, the total RNA count, cell cycle score and mitochondrial reads were regressed out. Principal-component analysis and UMAP with the first 20 dimensions was performed, followed by identifying clusters using a resolution of 0.20. Cell types associated with clusters were identified using established stromal markers^10,69^ and epithelial cell markers specific to liver and pancreas^36,70,71^.

#### Cluster Signatures for Single Cells

Network-based normalization of cluster signatures amongst single cells was achieved by first identifying a subset of genes whose expression was strongly correlated with its own cluster assignment (r > 0.4). For the PDX models, using the subset of highly correlated genes per cluster, if more than 30% of the cluster was captured per single cell, we calculated a centroid score by taking the mean of normalized UMI count. For patient CNB samples, centroid was calculated for all single cells if more than 20% of the cluster was captured. The percentage of cluster genes expressed within each cell type was calculated by taking the mean number of genes with UMI count above 0 per cluster across all the single cells within each cell type. Single cells were grouped into classical, quasibasal and basal based on the C1vC23 differential signature, classified as basal if the differential signature was < −0.1, quasibasal if they are −0.1 to 0.1 and classical if they were > 0.1.

#### Biomarker Prioritization

Cluster 1 and 23 specific networks were generated using Cytoscape 3.7.2 ^55^, where we calculated the closeness centrality of each gene to its respective cluster. Gene to cluster correlation was calculated by computing the Pearson correlation between each gene and its corresponding cluster score across our 48 PDX models. First degree nodes (FDN) were identified as directly connected nodes to CRISPR associated vulnerabilities *DNM2* and *MAPK11*.

The R packages ggplot2^72^ and ComplexHeatmap^52^ were used to plot the comparison between the gene to cluster correlation and closeness centrality and the gene expression heatmap of first-degree nodes, respectively.

#### Multiplex IHC-IF staining and data analysis

Formalin fixed and paraffin embedded (FFPE) PATX samples were sectioned into 3 μm thick sections and placed on positive charged slides. Sections were deparaffinised by baking at 60 °C for 1 hour, then rehydrated by serial passage thorough xylene and graded alcohol. All sections were subjected to an initial heat-induced epitope retrieval (HIER) in 10 mM citrate buffer with 0.05% Tween20, pH 6.0, at 95 °C for 15 minutes using a BioGenex EZ retrieval microwave. Subsequent HIER for Opal development was done using fresh citrate buffer at 95 °C for 10 minutes. All sections were initially blocked for endogenous peroxidase using Bloxal (Vector Labs SP6000). After and before each primary incubation, sections were blocked using 2.5% serum (Vector Labs S1012). Opal, indirect, and direct immunofluorescence methods were used. First node protein targets were developed using Opal methods, VSIG1 (Thermofisher MAB 4818, at 1/2000), and Vimentin (Cell Signaling Technologies 5741, at 1/1600). Ki67 (Cell Signaling Technologies 9129, at 1/400) was developed indirectly using anti-rabbit secondary, Alexa 680 (Thermofisher A32802, 1/500). Lastly, HLA conjugated to Alexa 647 (Abcam 199837, at 1/1000) was used to aid in tissue segmentation. Sections were then counterstained with DAPI.

Slides were imaged using Vectra 3.0 Automated Quantitative Pathology Imaging System (Akoya Biosciences). Image processing and analysis was performed using inForm Software v2.4 (Akoya Biosciences). For a subset of images from each PATX, the following was performed: Images were unmixed and autofluorescence was removed. Then, tissue was segmented as tumor, stroma or other based on training regions and pattern recognition of DAPI and HLA stain. This was followed by cell segmentation using DAPI and HLA to segment nuclei, cytosol, and membrane. Phenotyping was performed for each marker individually by selecting representative positives for algorithm training and allowing the software to select the rest. Batch analysis of all images was performed using the segmentation and phenotyping algorithm described above. At this threshold of detection, VSIG+/VIM+ cells represented a negligible population.

Spatial and data analysis was performed using phenoptrReports (Akoya Biosciences), an R script package. Briefly, all single cell phenotype data was merged, aggregated and consolidated for each marker. Consolidated data was analyzed based on the phenotypes of interest. Using the XY coordinates of each cell, spatial relationships between cell types was visualized using the phenotrReports GUI. All data was graphed using GraphPad Prism version 8.0.0 for Windows, GraphPad Software, San Diego, California USA, www.graphpad.com.

#### Feature Barcoding Vector

The feature barcode vector (LentiCRISPR-E-10xcs1) was built using the pLentiCRISPR-v2 (addgene: 52961) as the base vector. All of the molecular modifications were performed by Epoch Life Sciences (Missouri City, Tx). The pLentiCRISPR-v2 was modified to an optimized sgRNA scaffold^73^ that included the 3’ 10x capture sequence 1: (cgtttCagagctaTCGTGgaaaCAGCAtagcaagttaaaataaggctagtccgttatcaacttgaaaaagtggcaccgagtcggtgcGCTTTAAGGCCGGTCCTAGCAAtttttt); a ccdb bacterial expression cassette between the BsmBI restriction sites was introduced to reduce background during sgRNA cloning and library generation; and an N-terminal Flag-sv40 NLS was added to Sp. Cas9-nucleoplasmin NLS-P2A-Puro.

#### Preparation of Feature Barcoded sgRNA Knockout Populations

Lentiviral transductions of four separate feature barcode sgRNA vectors (targeting *ABCG8*, *ILK*, *SMAD4* and *ZEB1*), were conducted on separate cell populations for both the PATC69 and PATC53 PDX lines. Lentivirus was concentrated through ultra-centrifugation, resuspended in 200 μL of PBS, and stored at −80 °C until use. For each condition, 1×10^6^ cells were transduced in 10 cm treated plates (Corning) using 8 μg/mL Polybrene (Sigma-Aldrich). Media was replaced after a 16-hour incubation, and each cell population was washed with PBS and then placed under puromycin selection for 72 hours. Following selection, all conditions were cultured for 22 days (or 10 days post “CRISPR Screen Injection point”) to match the in vitro CRISPR screen control-separation profile at day 10 (Extended Data Figure 2A). Prior to library preparation for scRNAseq, knockout populations were combined in equal proportion for each PDX line. Library preparation was conducted on a total of 10,000 cells per PDX line, resulting in an approximate coverage of 2500 cells per condition.

#### sgRNA Phenotype Confirmation and Confirmation of Site-Specific Cutting

Utilizing the same feature barcoded populations prepared for scRNAseq, 1500 cells/well were seeded in triplicate in 12 well tissue culture plates (Corning) immediately following 72 hours of 2 μg/mL Puromycin selection. Cells were then cultured for a minimum of 10 doublings. Individual plates were then stained with 0.5% crystal violet (in 25% methanol) for 2 hours. Plates were washed in water, dried overnight and then digitally scanned. After digitally scanning the plates, crystal violet was dissolved in equal volumes of 1% SDS, and 200 μL of each sample was moved into 96-well plates to measure absorbance at 570 nm. Relative growth was quantified based on the internal sg*ABCG8* negative control. All data was graphed using GraphPad Prism v 8.0.

Puromycin selected cells were collected for Sanger sequencing to confirm sgRNA induced indel formation relative to non-infected populations. Cell pellets for each sgRNA across both PATC69 and PATC53, 1×10^6^ cells each, were isolated at Day 12 and Day 40 for each PDX line (sgRPS27A indels representative at Day 12, all other sgRNAs at Day 40). Cell pellets were centrifuged, washed once with PBS, and frozen at −80 °C. All Sanger sequenced regions were normalized against respective non-transduced PDX line populations, 1×10^6^ cells/pellet. Primers for each cut site were developed to allow for 400 - 800 bp products, and primer sites were run on 2% agarose gels and extracted following amplification. Site-specific sequencing primers were utilized for Sanger sequencing, and indels distributions were calculated using the Synthego ICE Analysis Tool^74^.

#### Feature Barcoding Analysis

Feature barcoded scRNAseq data were analyzed using Seurat3.1^68^. Each cell line was individually evaluated. Cells expressing more than 350 genes and less than 25% mitochondrial reads were retained and subsequently log normalized, variant stabilized, and total RNA count, mitochondrial reads and cell cycle were regressed out. All guide-level samples for each PDX line were merged using the Seurat Anchor Cell feature to provide a direct point of comparison for PATC69 and PATC53 perturbations. A total of 10,113 PATC69 (3449 *ABCG8*, 1962 *ILK*, 2662 *SMAD4*, and 2040 *ZEB1* knockout cells) and 11,439 PATC53 (3414 *ABCG8*, 2952 *ILK*, 2819 *SMAD4*, and 2254 *ZEB1* knockout cells) cells were retained and analyzed for processing. Principal-component analysis and UMAP with the first 20 dimensions was performed, with clustering performed at 0.15 resolution for. Cluster centroids were calculated using the method described above for the patient CNB samples.

## ACKNOWLEDGMENTS

We thank the Advanced Technology Genomics Core (ATGS), the UTMDACC Flow Cytometry and Cellular Imaging Core Facility, the UTMDACC Department of Veterinary Medicine, all funded by a Cancer Center Support Grant (P30 CA016672). We thank David Aten and Jordan Pietz at the UTMDACC Medical Graphics and Photography Department funded by Cancer Center Support Grant (P30 CA016672). We thank the UTMDACC Science Park NGS Facility, funded by CPRIT Core Facility Support Grants (RP120348 and RP170002). We thank Sisi Gao for her contributions editing the manuscript. We thank our colleagues at the Institute of Applied Cancer Science (IACS), Justin Huang, Stephanie Schmidt, and Vandhana Ramamoorthy, for technical assistance and suggestions. We thank David Pollock at ATGS for technical assistance and advice. G.F.D. was supported by the Sewell Family Endowed Chairmanship in Genomic Medicine, NIH/NCI P01 CA117969 12, and the UT MD Anderson Cancer Moon Shots Program. S.S was supported by the CPRIT Research Training Grant (RP170067). J.R. and I.H. received support from the Paula-Altman Goldstein Discovery Fellowship. J.R. also received support from P01 CA117969 12. W.Y. is supported by the PanCAN-AACR Pathway to Leadership Grant (16-70-25-YAO) and the Pancreatic Cancer Action Network Translational Research Grant (19-65-YAO). TH is a CPRIT Scholar in Cancer Research, and is supported by NIGMS grant R35GM130119 and MD Anderson Cancer Center Support Grant P30 CA016672. TH is a consultant for Repare Therapeutics.

## AUTHOR CONTRIBUTIONS

J.L.R., S. Srinivasan., A.C. and G.F.D. conceived and designed the study. J.L.R., W.Y., M.P., A. Machado. performed the experiments. S. Srinivasan, S. Seth, E.K. and C.A.B analyzed the data. PDX generation was guided by the lab of M.K. Single-cell RNAseq data of PDXs were provided by C.Y.L., I.H. and A.V. Single-cell RNAseq data from patients were provided by J.J.L., P.A.G., and A. Maitra. Clinical annotations for PDXs were provided by M.S. and E.J.K. Guidance on experimental and computational aspects of the study was provided by W.Y., J.R.D., C.A.B., G.G., A.V., T.P.H., A. Maitra, and T.H. The project was led and supervised by A.C. and G.F.D. The manuscript was written by J.L.R., S. Srinivasan, A.C. and G.F.D. and edited by A.K.D., A. Maitra and T.H. All authors approved the final manuscript.

