## Extended Figures and Legends for "Diversity Across the Pancreatic Ductal Adenocarcinoma Disease Spectrum Revealed by Network-Anchored Functional Genomics"

**EXTENDED DATA FIGURE LEGENDS:**

**Extended Data Figure 1: Characterization of the PDAC PDX cohort and subsequent refinement and annotation of the co-expression network (related to Fig. 1).** (**a**) Moffitt classification of PDAC PDX models (n=48) using the PDX-specific gene set. Consensus clustering of normalized gene expression was performed to classify models into classical (blue top annotation) and basal-like (yellow top annotation). (**b**) Dot plot demonstrating the change in the lowest gene-pair pearson correlation per Log Likelihood Score (lls) cutoff for inclusion in the network. (**c**) Results from the UNCOVER analysis reveal complementary gene mutations associated with cluster centroid-based enrichment. (**d**) Combined mutations in *ACVR2A*, *RREB1* and *MARK2* showed statistically significant association with Cluster 21 enrichment ($p < 0.05$). (**e**) Treemap display of GO-annotated pathways of the PDAC co-expression network on a cluster by cluster basis.

**Extended Data Figure 2: RNAi screening of PDAC-prioritized surface proteins in PDX models initially informs on a vulnerability diversity within the PDAC cohort (related to Fig. 2).** (**a**) Schematic of parallel *in vitro* and *in vivo* RNAi- and CRISPR-based screening strategies. Day 1 of screening for RNAi started immediately following 72 hours of puromycin selection, while day 1 of CRISPR screening started at 12 days post-transduction, for both *in vitro and in vivo* time-points, to allow for initial sgRNA cutting prior to *in vivo* implantation into the mouse pancreas. In both cases, the Reference population for determining barcode dropout was based on barcode distribution immediately following puromycin selection. (**b**) Outline detailing which PDX line was used for RNAi- and CRISPR-based screening, and the Moffitt classification for each model used. (**c**) Common PDAC-associated mutations for the four PDX lines used for genetic screening. (**d - g**) Comparative dot plots highlighting parallel gene-level p-values, generated by RSA, for Day 30 of the *in vitro* and *in vivo* PATC69 (d), PATC124 (e), PATC53 (f), and PATC153 (g) screens. (**h**) Venn diagram of significant ($p < 0.05$) RNAi-defined gene targets, identified *in vivo*. Generalized protein function is highlighted for each displayed target.

**Extended Data Figure 3: Low-fat BAGEL for optimized analysis of small custom CRISPR libraries and *in vivo* functional genomics (related to Figure 2). (a)** Kernel density estimate plots comparing the log2 fold-change distribution of all essential and non-essential guides (n=285, n=280 respectively) for PATC69 *in vitro* Day 12 screen demonstrating the differences in dropout between the two distributions. **(b)** Gene-wise BFs of essential and non-essential genes from PATC69 *in vitro* Day 12 screen calculated from BAGEL v1 designed for genome-wide CRISPR screens. Gene-wise BF is calculated as a sum of guide level BFs. **(c)** Gene-wise Bayes Factors of essential and non-essential genes PATC69 *in vitro* Day 12 screen calculated from Low-fat BAGEL designed for custom library CRISPR screens. Gene-wise BF is calculated as a sum of guide-level BFs. **(d)** Precision-Recall of BFs of essential and non-essential genes from the PATC69 *in vitro* Day 12 screen comparing the performance of BAGEL v1 and Low-fat BAGEL. **(e)** Comparison of F measures of known genome-wide CRISPR screen analytical methods to Low-Fat BAGEL across PATC69 and PATC124 *in vitro* and *in vivo* screens.

**Extended Data Figure 4: Functional genomics quality control for RNAi- and Low-fat Bagel-analyzed CRISPR screening datasets (related to Fig. 2).** (**a**) Size-normalized read count distribution for each *in vitro* and *in vivo* RNAi screen. (**b**) Fold-change separation comparison for shRNAs (n =10) targeting negative control luciferase (non-targeting) vs. positive control genes *PSMA4* and *RPL30*. (**c**) CPM-normalized read distribution for each *in vitro* and *in vivo* CRISPR screen. (**d**) Density plots of essential and nonessential guide log2 fold change dropout for PATC69. An average of three replicates per time point is plotted. **(e)** Combined precision-recall curves for PATC69 screens using Low-fat BAGEL. **(f)** Density plots of essential and non-essential guide log2 fold change dropout for PATC124. An average of three replicates per time point is plotted. **(g)** Combined precision-recall curves for PATC124 screens using Low-fat BAGEL. **(g)** Density plots of essential and non-essential guide log2 fold change dropout for PATC69. An average of three replicates per time point is plotted. **(h)** Combined precision-recall curves for PATC69 screens using Low-Fat BAGEL.

**Extended Data Figure 5: Parallel *in vitro* CRISPR characterization of PDX models identifies functional diversity within the PDAC cohort (related to Fig. 2).** (**a**) Venn diagram of functional targets derived from *in vivo*, orthotopically implanted, CRISPR screening of PDX lines using a quantile-normalized BF > 1. (**b - d**) *in vitro* CRISPR screening results from PATC69 (b), PATC124 (c), and PATC53 (d) PDX lines overlaid onto a merged PPI force-directed diagram. The BF of each gene indicates the degree of vulnerability of the gene. (**e - g**) Network anchoring of *in vivo* functional dependencies, represented as quantile-normalized BFs, to inform on cluster-associated vulnerability context in PATC69 (f) PATC124 (g), and PATC53 (h) PDX lines.

**Extended Data Figure 6: Single-cell subpopulation characterization and marker gene selection for patient core biopsies (related to Fig. 4).** (**a**) UMAPs of 25,954 single cells separated by the seven patient core needle biopsies. Analysis was performed using Seurat 3.1 and clusters were identified using a resolution of 0.2. (**b**) Expression of known marker genes was used to identify cell type of each UMAP-identified cluster.

**Extended Data Figure 7: Selection of sgRNAs and validation of site-directed cutting of C1vC23-associated genetic targets (related to Fig. 6).** (**a**) sgRNAs used for CRISPR knockout and feature-barcode tracking scRNAseq. Guides were selected based on dropout activity in the parallel *in vitro* and *in vivo* CRISPR screens for PATC69 and PATC53. (**b - k**) Indel frequencies and patterns for selected sgRNAs in both PATC69 and PAYC53. Indel frequency was calculated using Sanger sequencing and the Synthego ICE CRISPR analysis tool (see Methods).

**Extended Data Figure 8: Colony formation following sgRNA transduction and intratumoral C1vC23 characterization of sequenced PDX populations (related to Fig. 6).** (**a** - **b**) Crystal violet staining of colony growth for PATC69 [left panel] and PATC53 [right panel]. Colonies were seeded immediately following puromycin selection (1,500 cells/well), and cultured for a minimum 10 doublings to confirm dropout activity relative to the sgABCG8 negative control. (**c - d**) UMAPs detailing the C1vC23 signature in each single-cell RNA sequenced for PATC69 [left panel] and PATC53 [right panel] *ABCG8* knockout populations.

**Extended Data Figure 1:** Characterization of the PDAC PDX cohort and subsequent refinement and annotation of the co-expression network (related to Fig. 1).

A

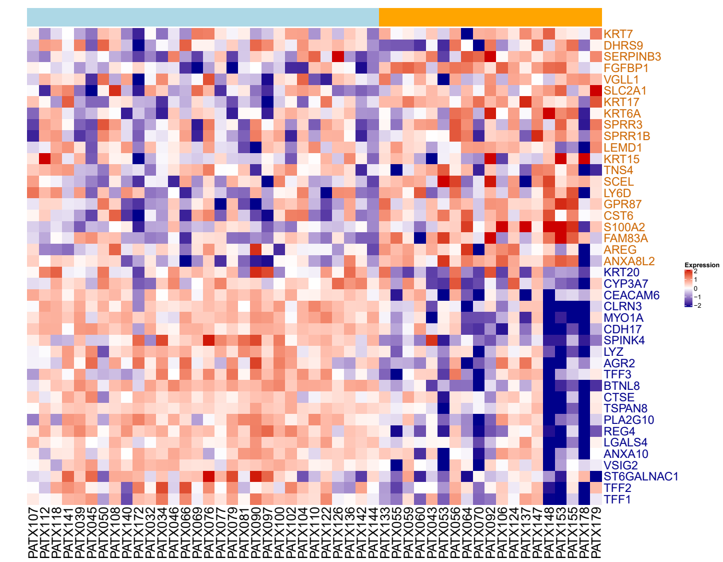


B

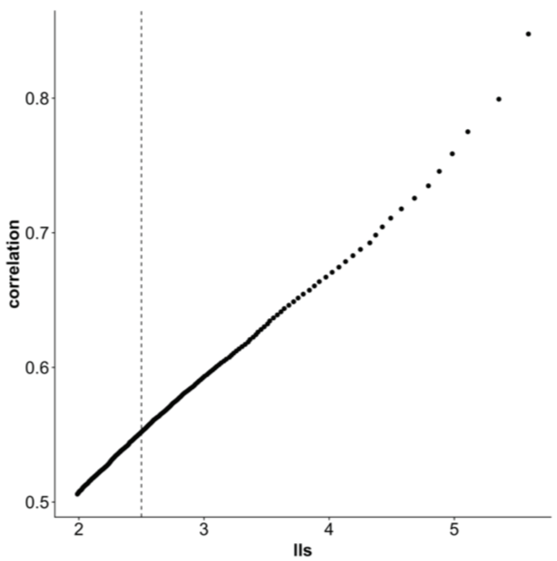


C

| Cluster | Mutation 1 | Mutation 2 | Mutation 3 | OFC | p-value |
|---|---|---|---|---|---|
| cluster_1 | SETD2 | PLXNB2 | SIN3B | 3.526925 | 0.752475 |
| cluster_2 | KDM6A | BCORL1 | BRCA2 | 5.498805 | 0.277228 |
| cluster_3 | KDM6A | BCORL1 | SDK1 | 4.765432 | 0.485149 |
| cluster_4 | NF1 | BRCA2 | RREB1 | 5.238452 | 0.326733 |
| cluster_5 | RREB1 | MARK2 | ACVR2A | 6.091605 | 0.217822 |
| cluster_6 | SDK1 | NF1 | PALB2 | 5.14112 | 0.267327 |
| cluster_7 | RREB1 | RNF43 | MARK2 | 5.484781 | 0.287129 |
| cluster_8 | SDK1 | ACVR2A | MARK2 | 5.674092 | 0.168317 |
| cluster_9 | RREB1 | BRCA2 | RNF43 | 5.141292 | 0.445545 |
| cluster_10 | PLXNB2 | RNF43 | RREB1 | 5.237245 | 0.376238 |
| cluster_11 | SPTB | PALB2 | SDK1 | 5.575874 | 0.089109 |
| cluster_12 | SPTB | ITPR3 | BRCA2 | 4.064193 | 0.782178 |
| cluster_13 | PALB2 | ITPR3 | SIN3B | 4.180341 | 0.653465 |
| cluster_14 | PLXNB2 | PALB2 | ITPR3 | 4.296772 | 0.712871 |
| cluster_15 | SDK1 | PLXNB2 | RREB1 | 4.013054 | 0.792079 |
| cluster_16 | ITPR3 | SETD2 | SMARCA4 | 3.520863 | 0.980198 |
| cluster_17 | PALB2 | TLE4 | ITPR3 | 4.769161 | 0.574257 |
| cluster_18 | ACVR2A | ITPR3 | BRCA1 | 5.513736 | 0.415842 |
| cluster_19 | PALB2 | CALD1 | SMARCA4 | 6.269093 | 0.287129 |
| cluster_20 | SDK1 | CALD1 | AKT2 | 4.720414 | 0.653465 |
| cluster_21 | ACVR2A | RREB1 | MARK2 | 7.206207 | 0.029703 |
| cluster_22 | SDK1 | PALB2 | CALD1 | 4.122304 | 0.871287 |
| cluster_23 | CALD1 | SMARCA4 | PALB2 | 4.131839 | 0.920792 |
| cluster_24 | ACVR2A | TP53BP2 | CALD1 | 4.385642 | 0.831683 |
| cluster_25 | PALB2 | SETD2 | CALD1 | 4.767625 | 0.524752 |
| cluster_26 | SMARCA4 | RREB1 | PALB2 | 4.372286 | 0.80198 |
| cluster_27 | CALD1 | AKT2 | BCORL1 | 4.055904 | 0.792079 |
| cluster_28 | AKT2 | PLXNB2 | CALD1 | 4.196129 | 0.772277 |
| cluster_29 | RNF43 | PLXNB2 | PALB2 | 4.640496 | 0.366337 |
| cluster_30 | PALB2 | CALD1 | PLXNB2 | 5.071861 | 0.60396 |
| cluster_31 | PALB2 | SMARCA4 | CALD1 | 4.305349 | 0.881188 |

D

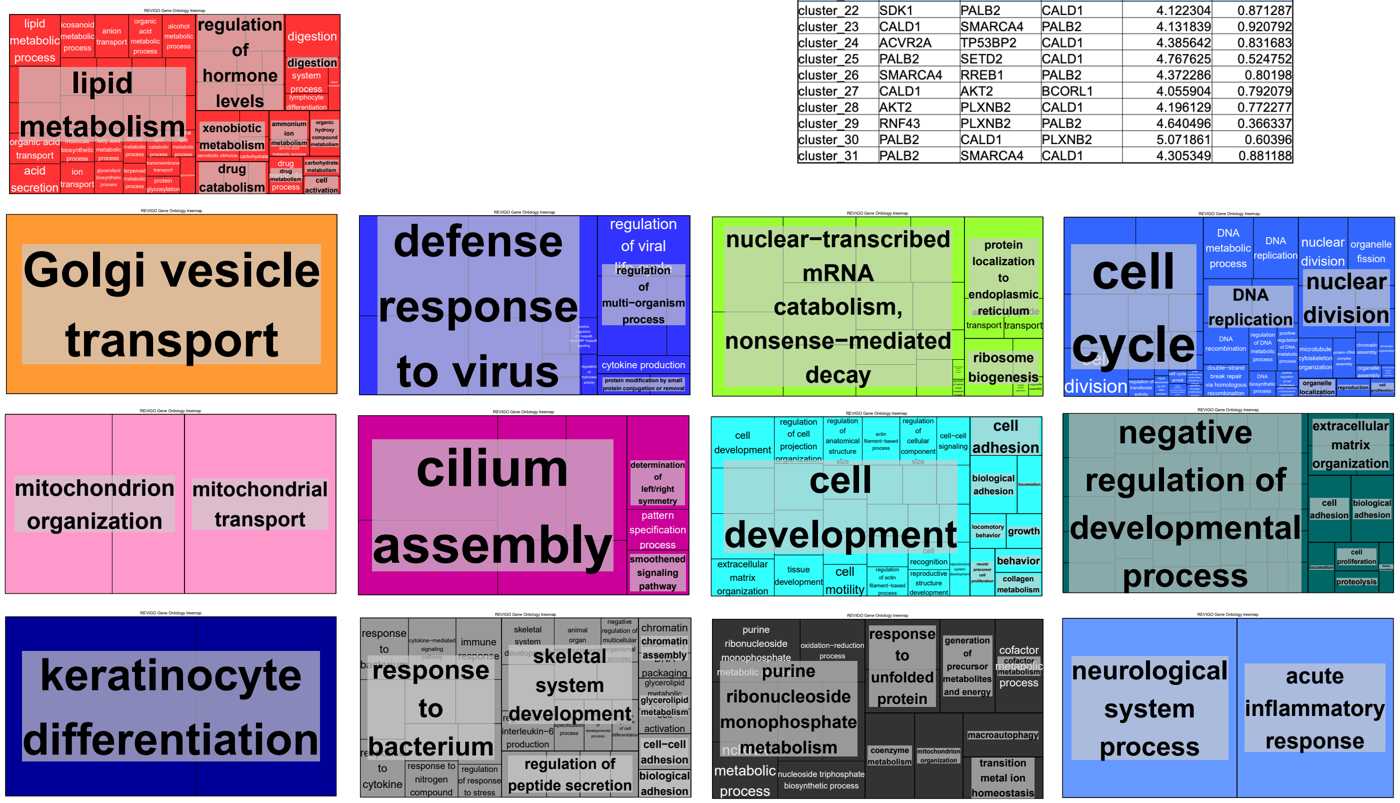

**Extended Data Figure 2:** RNAi screening of PDAC-prioritized surface proteins in PDX models initially informs on a vulnerability diversity within the PDAC cohort (related to Fig.2)

A

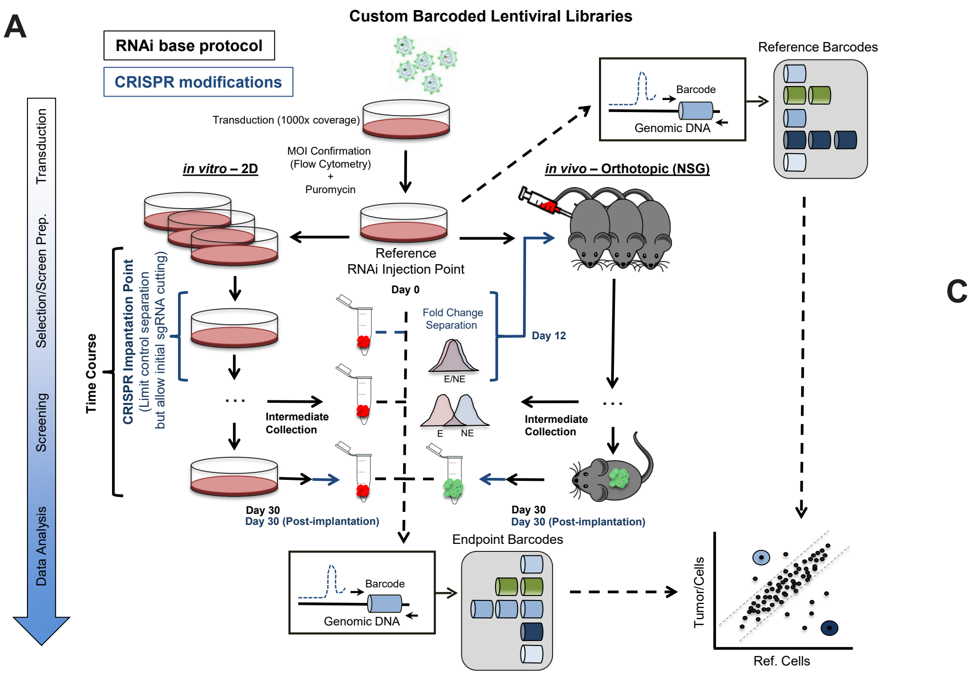


B

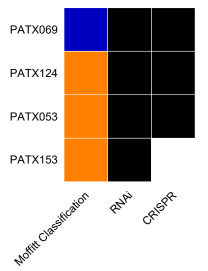


C

| Model | Gene | Variant effect | AA_Change | AF |
|---|---|---|---|---|
| PATC69 | CDKN2A | FRAME_SHIFT | p.Arg156fs | 1.00 |
| | ATM | NON_SYNONYMOUS_CODING | p.Phe2732Val | 1.00 |
| | KRAS | NON_SYNONYMOUS_CODING | p.Gly12Asp | 1.00 |
| | TP53 | CODON_INSERTION | p.Asn235dup | 1.00 |
| | PALB2 | NON_SYNONYMOUS_CODING | p.Asp219Gly | 0.40 |
| | BRAF | NON_SYNONYMOUS_CODING | p.Cys95Ser | 0.33 |
| PATC124 | ARID1B | NON_SYNONYMOUS_CODING | p.Gly168Asp | 1.00 |
| | TP53 | FRAME_SHIFT | p.Arg333fs | 1.00 |
| | BRCA1 | FRAME_SHIFT | p.Ser1676fs | 1.00 |
| | SMAD4 | SPLCE_SITE_DONOR+INTRON | NA | 1.00 |
| | ATM | NON_SYNONYMOUS_CODING | p.Ser1104Cys | 0.73 |
| | KRAS | NON_SYNONYMOUS_CODING | p.Gly12Asp | 0.67 |
| PATC53 | TP53 | STOP_GAINED | p.Arg306* | 1.00 |
| | KMT2D | NON_SYONYMOUS_CODING | p.Prp2557Leu | 1.00 |
| | ARID1A | FRAME_SHIFT | p.Ser186fs | 1.00 |
| | ARID1A | NON_SYONYMOUS_CODING | p.Pro158Ser | 0.98 |
| | ALK | NON_SYONYMOUS_CODING | p.Ala1601Thr | 0.74 |
| | KRAS | NON_SYONYMOUS_CODING | p.Gly12Asp | 0.72 |
| PATC153 | TP53 | NON_SYNONYMOUS_CODING | p.Arg175His | 1.00 |
| | AR | CODON_CHANGE/DELETION | p.Gly465_Gly4 | 1.00 |
| | PIK3CA | NON_SYNONYMOUS_CODING | P.Glu542Lys | 0.74 |
| | TET2 | NON_SYNONYMOUS_CODING | p.Try888His | 0.54 |
| | TET2 | NON_SYNONYMOUS_CODING | p.Pro1744Ser | 0.47 |

D

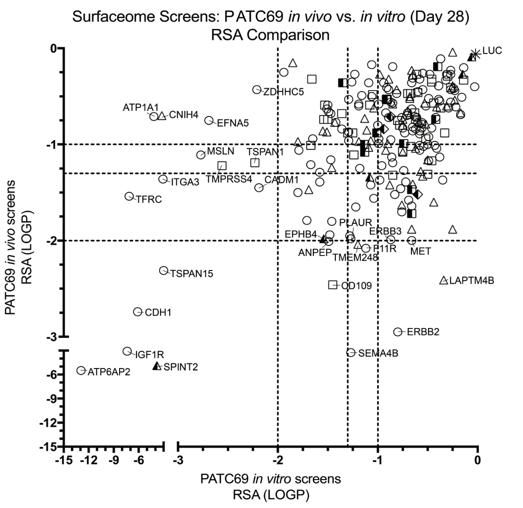


E

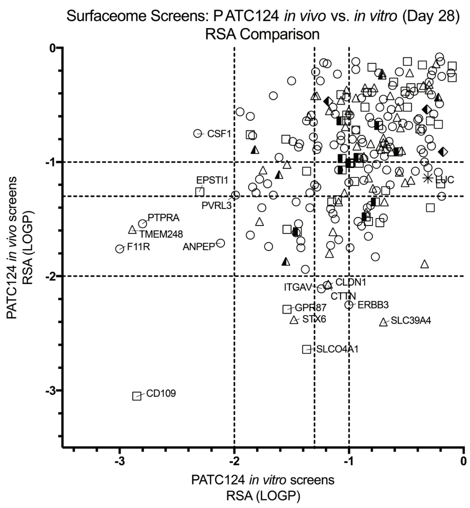


F

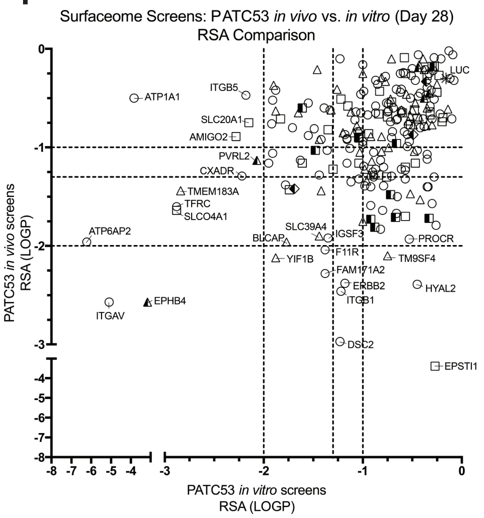


G

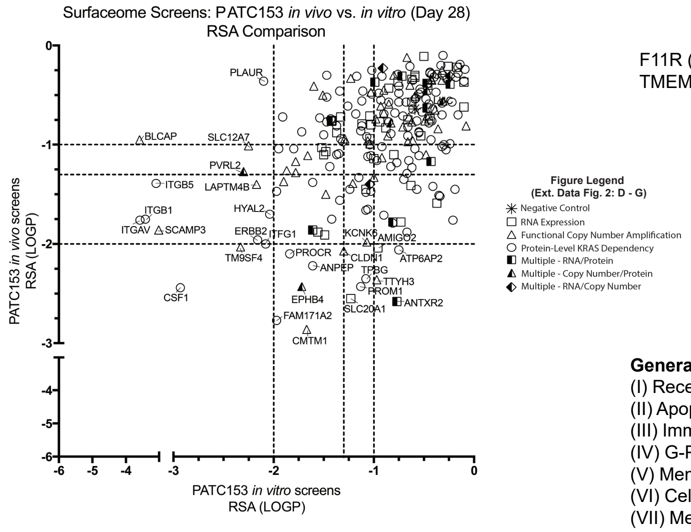


H

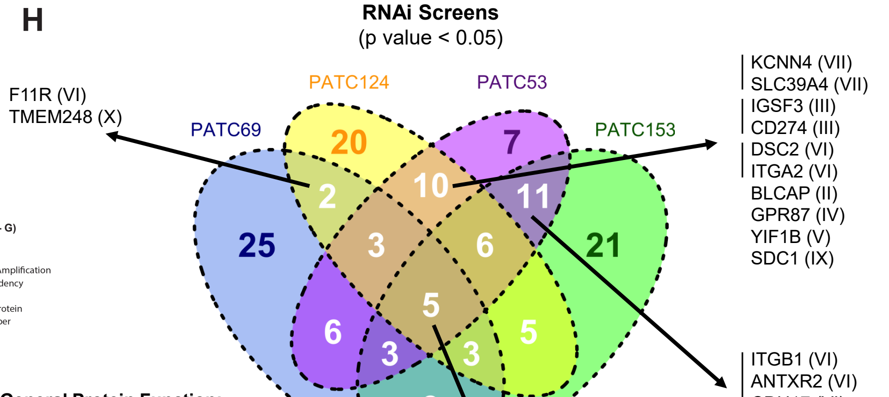


**General Protein Function:**
(I) Receptor Tyrosine Kinase
(II) Apoptosis
(III) Immunoregulatory
(IV) G-Protein coupled receptor
(V) Membrane trafficking Protein
(VI) Cell-cell junction/adhesion
(VII) Membrane channel/transport
(IX) Surface Regulation/Differentiation/Internalization
(X) Undefined transmembrane proteins

**Extended Data Figure 3:** Low-Fat BAGEL for optimized analysis of small custom CRISPR libraries and *in vivo* functional genomics (related to Figure 2).

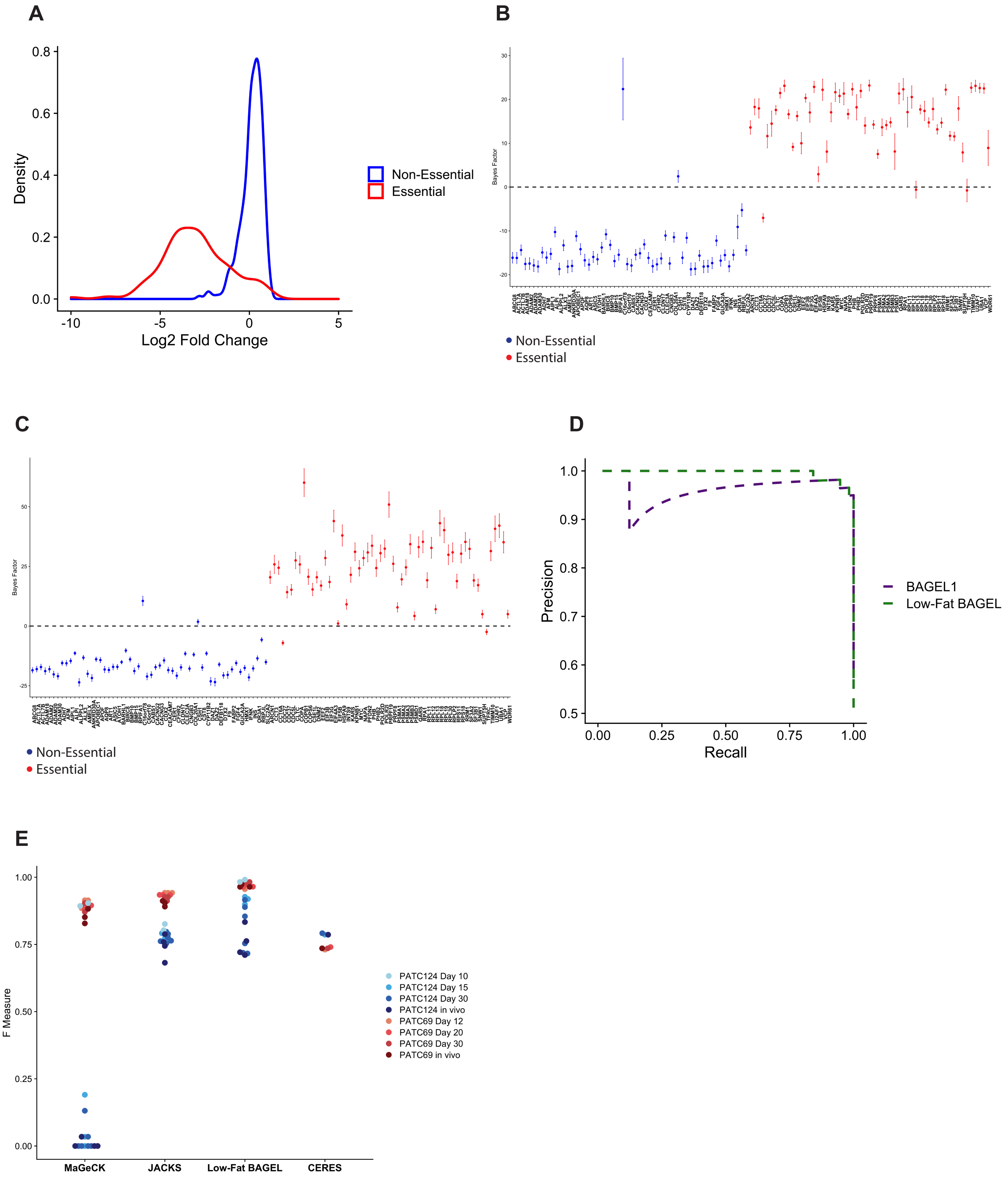

**Extended Data Figure 4:** Functional genomics quality control for RNAi and Low-fat Bagel analyzed CRISPR screening datasets (related to Fig. 2).

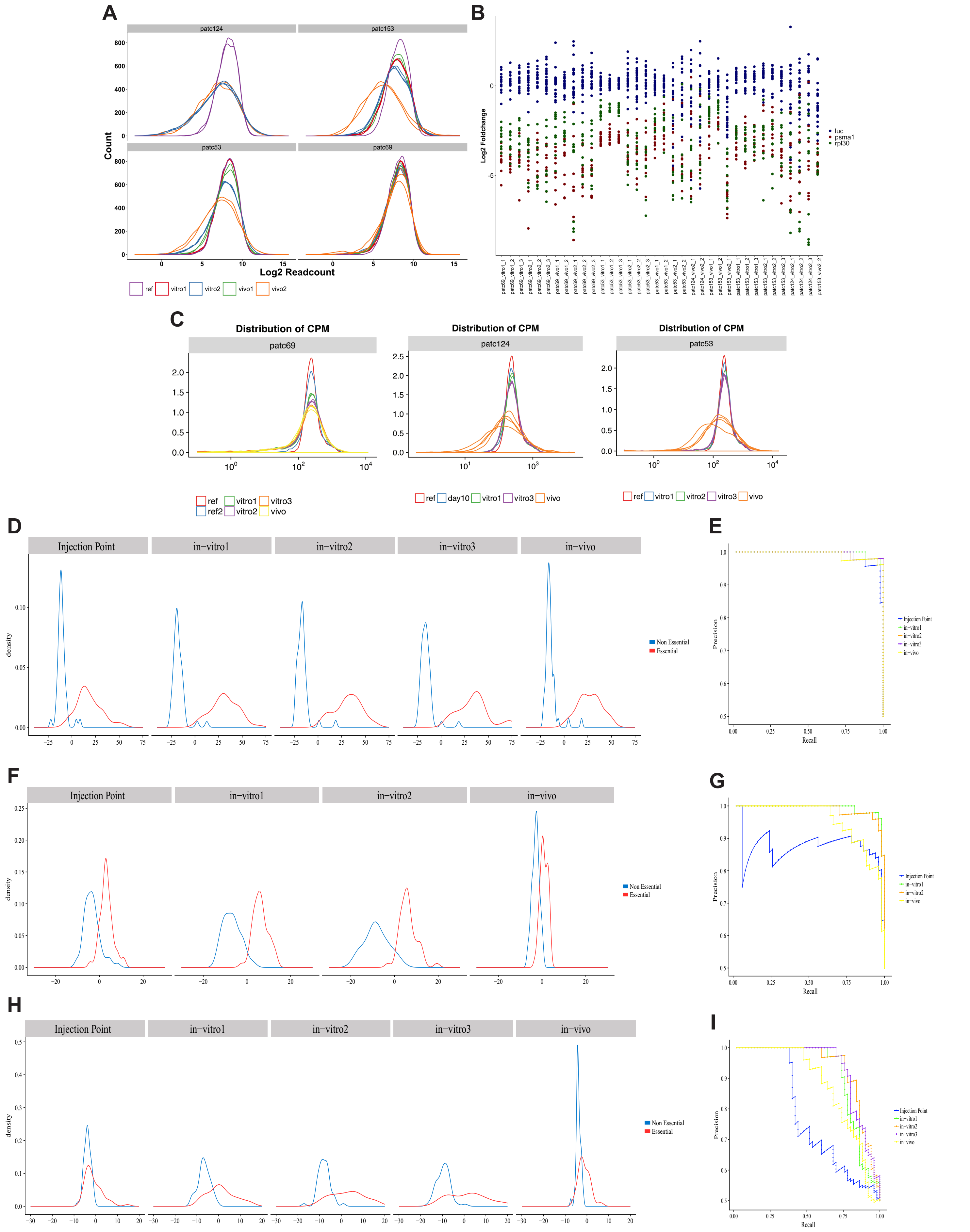

**Extended Data Figure 5:** Parallel *in vitro* CRISPR characterization of PDX models identifies functional diversity within the PDAC cohort (related to Fig. 2).

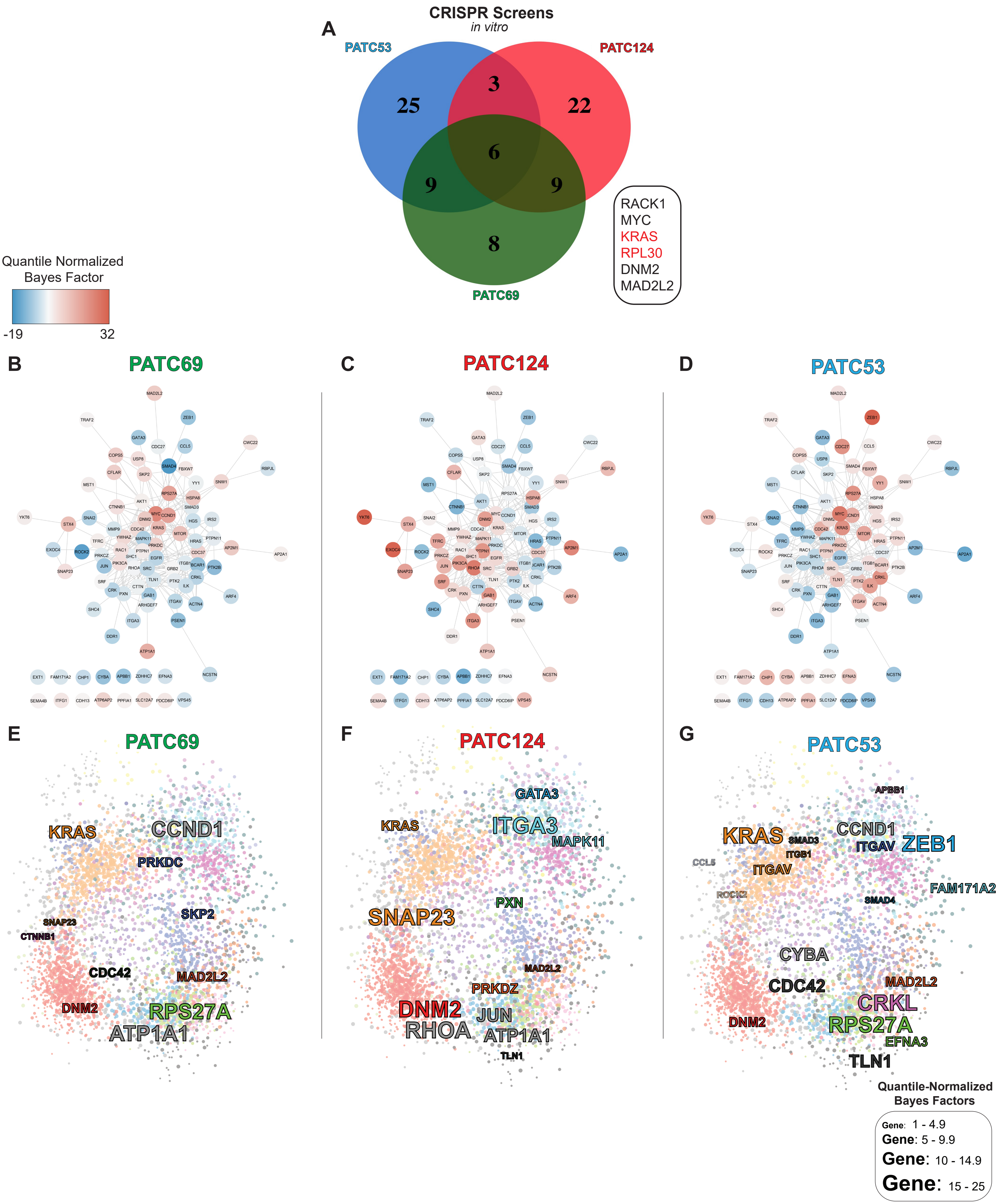

**Extended Data Figure 6:** Single-cell subpopulation characterization for PDAC patient core biopsies (related to Fig. 4).

**A**

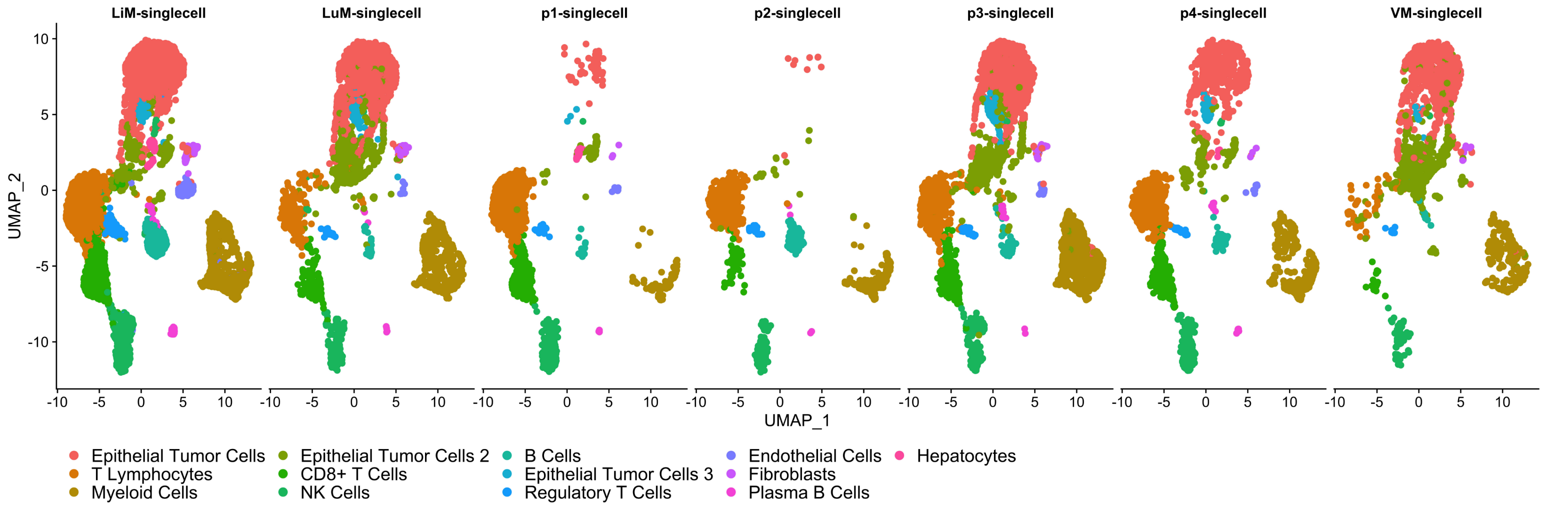


**B**

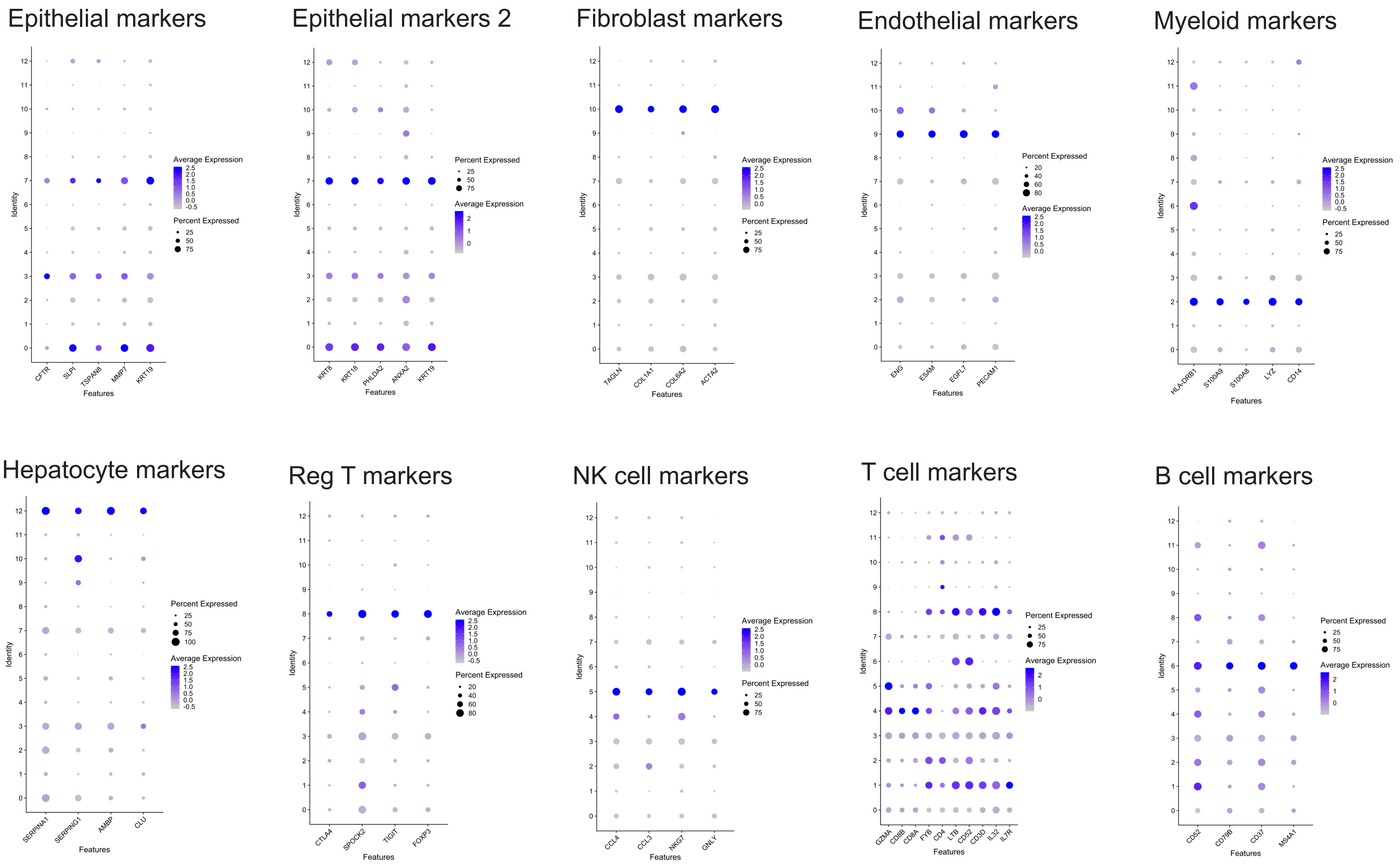

**Extended Data Figure 7:** Selection of sgRNAs and validation of site-directed cutting of C1vC23 associated genetic targets (related to Fig. 6).

A

| sgRNA | sgRNA Library ID | gRNA |
|---|---|---|
| sgABCG8_1 | pool_4\|sgABCG8_BB_2\|ABCG8 | CAGTGCGCTGACACCCGCGT |
| sgILK_1 | pool_4\|sgILK_BB_1\|ILK | ACATTGTAGAGGGATCCATA |
| sgSMAD4_1 | pool_4\|sgSMAD4_BB_1\|SMAD4 | GCAATGGAACACCAATACTC |
| sgZEB1_1 | pool_4\|sgZEB1_BB_2\|ZEB1 | CAGACCAGACAGTGTTACCA |
| sgRPS27A_1 | pool_4\|sgRPS27A_BB_2\|RPS27A | CCACCCGGCCCGTACCTCGA |

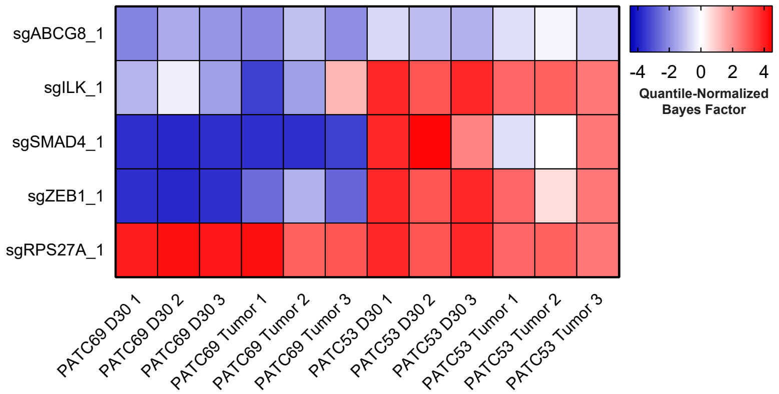


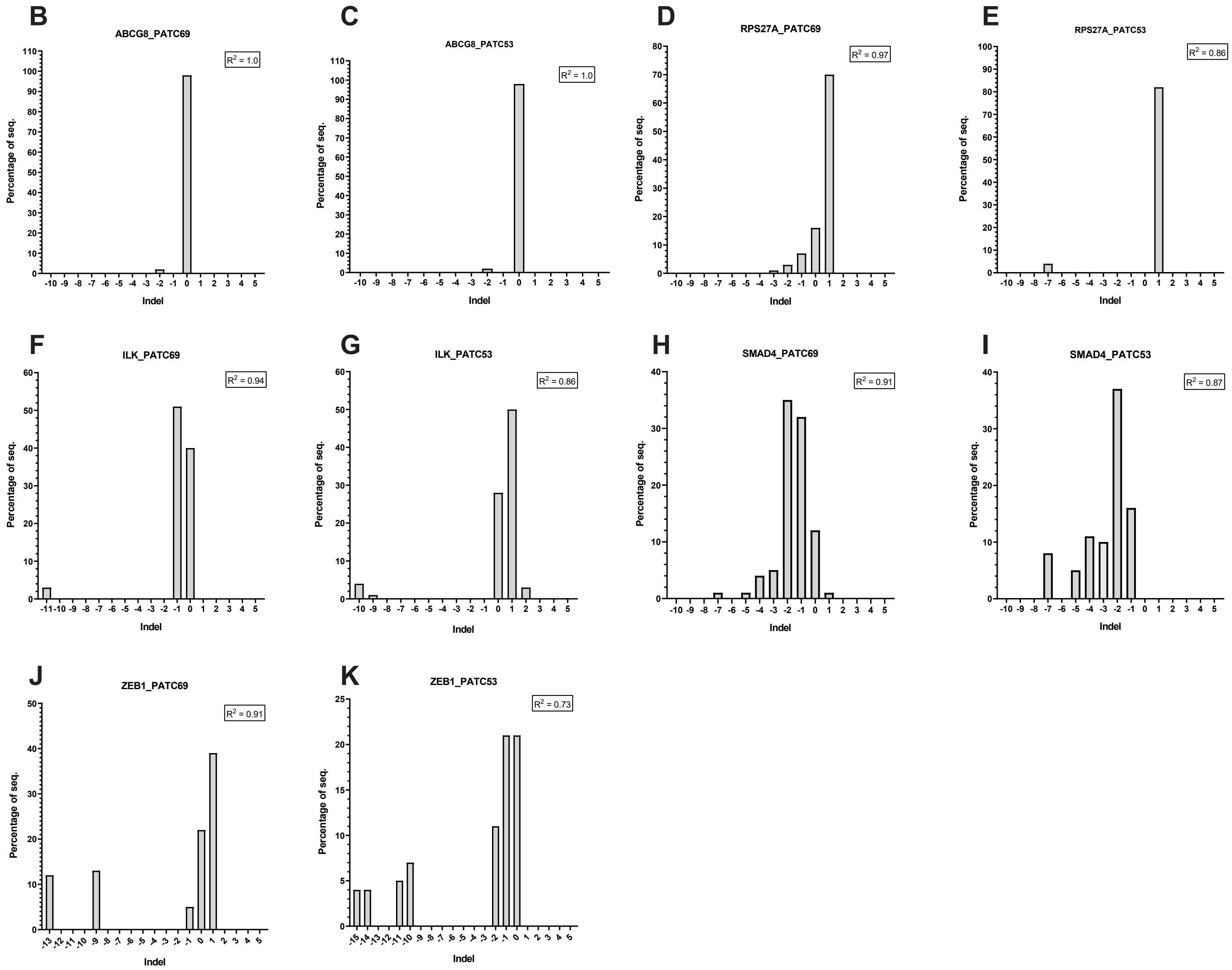

**Extended Figure 8:** Colony formation following sgRNA transduction and intratumoral C1vC23 characterization of sequenced PDX populations (related to Fig. 6).

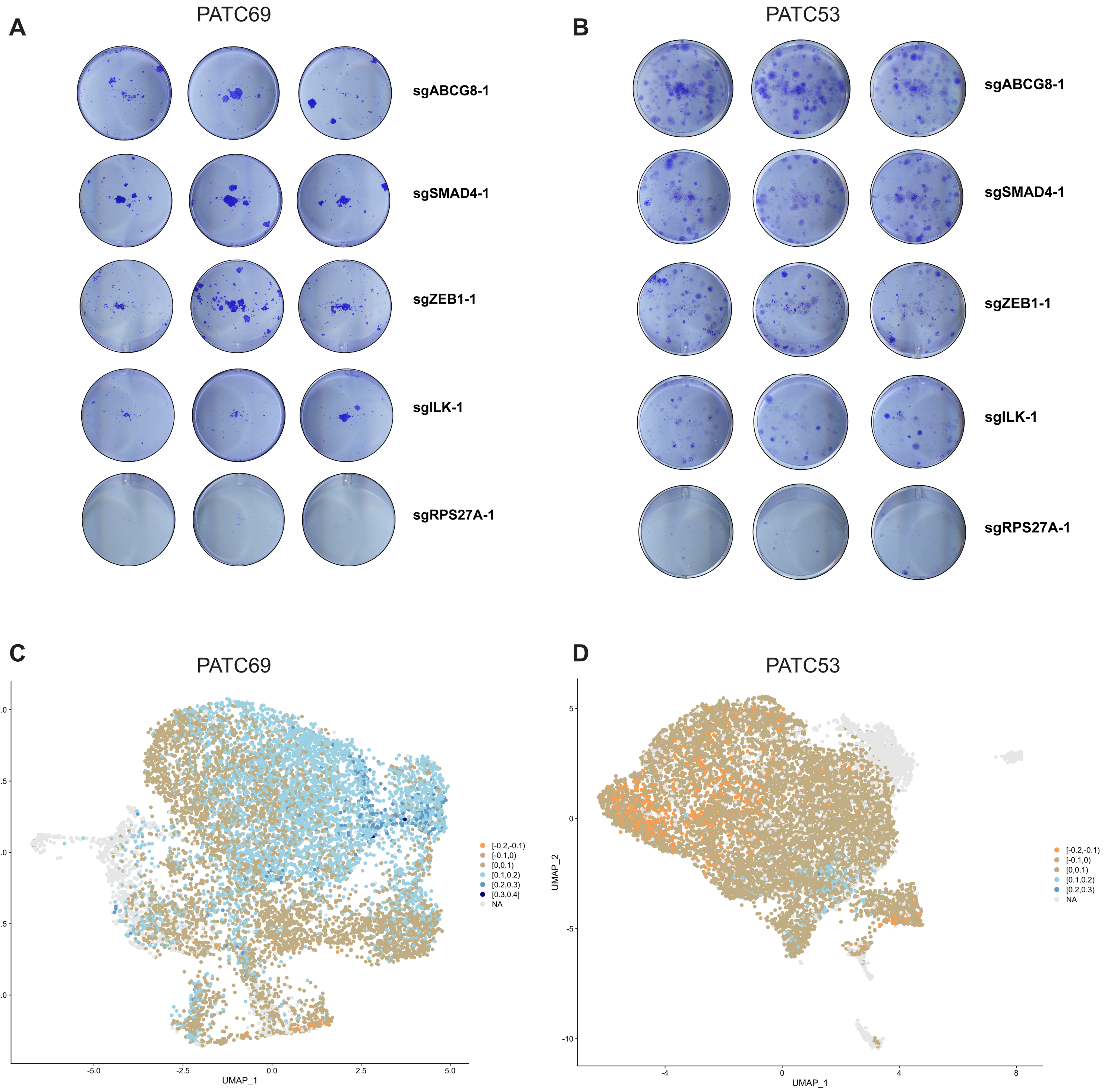
